# Phospotyrosine proteomics reveals novel Zap70 and Itk pathway targets downstream of TCR and CAR in Jurkat T cells

**DOI:** 10.1101/2025.11.03.686362

**Authors:** Aurora Callahan, Savannah S. Trychanh, Timothy Ro, Aisharja Mojumdar, Arthur R. Salomon

## Abstract

ζ-associated protein of 70 kDa (Zap70) and interleukin-2-inducible T cell kinase (Itk) propagate the primary and CD28-integrated phosphotyrosine (pY) signalling, respectively, to achieve full T cell activation. Despite their canonical roles in T cell activation, our understanding of how each kinase controls canonical and noncanonical pY signalling is incomplete. Here, using three T cell activation methods (soluble antibodies, APC-pMHC/TCR, and CD19-CAR/Raji), we evaluated the effects of two novel inhibitors, RDN2150 (RDN, Zap70) and Soquelitinib (Soq, Itk), on T cell activation. We validated the published working concentrations of RDN and Soq on phosphorylation of key T cell signalling proteins and on T cell activation markers, finding that RDN provides more complete inhibition of T cell signalling and activation. We used LC-MS/MS to evaluate how RDN and Soq treatment affected the phosphotyrosine (pY) signalling and proteome of T cells, finding that RDN, as opposed to Soq, completely downregulated the TCR signalling pathway. Finally, we identified new, noncanonical pY sites responsive to RDN and Soq, providing new insights into the pathways regulated by Zap70 and Itk. Together, our work provides a basis for further study on RDN and Soq, as well as a molecular roadmap for the effects of these inhibitors.

## 1 Introduction

T cell activation through the T cell receptor (TCR) is initiated by binding of a peptide-presenting major histocompatibility complex (pMHC) on an antigen-presenting cell. It requires the coordinated activity of tyrosine kinases, including the Src family kinase Leukocyte C-terminal Src kinase (Lck), the Syk family kinase ζ-associated protein of 70 kDa (Zap70), and the Tec family kinase interleukin 2 inducible T cell kinase (Itk)^1^. TCR-pMHC ligation promotes Lck^Y505^ dephosphorylation by CD45 and subsequent Lck^Y394^ autophosphorylation and phosphorylation of immunoreceptor tyrosine-based activation motifs (ITAMs) on TCR subunits CD3ɛ, CD3γ, CD3δ, and TCRζ, which Zap70 is then able to bind through its dual SH2 domains. Lck activates Zap70 and Itk through phosphorylation at Zap70^Y493^ and Itk^Y512^, respectively, and linker for activation of T cells (LAT), a scaffolding protein necessary for TCR signalling, is activated by Lck-mediated Zap70 phosphorylation at multiple tyrosine residues. Multiple scaffolding proteins, including SLP76-bound GADS and phospholipase C γ1 (PLCγ1), dock on LAT, allowing for the phase separation of the TCR signalosome and bringing substrates into proximity with Lck, Zap70, and Itk for phosphorylation and activation. Components of the TCR signalosome then propagate signal to reorganise the actin cytoskeleton and activate transcription factors, promoting T cell activation and feedback regulation of the signalling input (reviewed extensively in Gaud *et al*. 2018). Notably, mutations to Zap70 are associated with autoimmune disorders and loss of Zap70 ablates T cell activation, while inhibition of Itk alters the fate of T cells and is associated with better prognosis in immunotherapeutic intervention in various diseases^2,3^.

Chimeric antigen receptors (CARs), a novel immunotherapy for treatment of autoimmune disorders and cancers, are composed of domains from T cell signalling proteins with the goal of engaging a disease-associated antigen and initiating an immune response^4,5^. Clinically available CAR designs include the intracellular domain of TCRζ, as well as costimulation using the intracellular domain of either CD28 or 4-1BB, two T cell costimulatory receptors that enhance T cell activation and memory T cell formation, respectively^6,7^. Despite hundreds of clinical trials evaluating the safety and efficacy of CARs in humans, only six are currently approved for use as a last resort for relapsed or refractory cancers of blood cell lineage^6^. CAR T cell therapy is often plagued by non-specific T cell activation, low antigen sensitivity, and T cell exhaustion, leading to variable patient responses, at worst resulting in no tumour clearance, cytokine release syndrome, and neurotoxicity resulting from enhanced immune activity in the brain^8^. Recent literature suggests that altered T cell activation is the result of aberrant engagement of the TCR signalling pathway, including constitutive docking of active Lck on CD28 costimulated CARs, a lack of recruitment of key TCR signalling proteins after antigen ligation on 4-1BB costimulated CARs, and differential engagement with the TCR signalosome^9–11^. Researchers have demonstrated that disrupting the binding site of Itk in the CD28 costimulatory domain (PRRP to ARRA) or the GADS binding site (YMNM to FMNM) can significantly enhance the function of second-generation CARs with a CD28 domain, thereby reducing constitutive and continuous CAR activation^12,13^. Currently, efforts to restrain CAR-induced toxicities have come from Jurkat high-throughput screens evaluating combinatorial CAR libraries for sustained T cell activation with low exhaustion markers and the rational design of new CARs, including dual-antigen split CARs, and “fourth-generation CARs” known as T cells redirected for universal cytokine-mediated killing (TRUCKs)^14–22^.

Even with advances in the functional understanding of CAR T cells, basic research evaluating how CARs influence T cell biology, particularly T cell activation, is lagging.

Two small-molecule inhibitors with distinct effects on T-cell signalling have recently been described: RDN2150 (RDN) targeting Zap70 and Soquelitinib (Soq) targeting Itk^23,24^. RDN covalently inhibits Zap70 and prevents phosphorylation of LAT^Y220^, thereby destabilising the LAT/SLP76 complex. At concentrations less than 400 nM, RDN reduces interferon-γ and interleukin-17A, and is effective in treating psoriasis in murine models by topical application^23^. Soq is a highly selective covalent inhibitor of Itk, thought to reduce phosphorylation of PLCγ1^Y783^ and thus reduce T cell activation strength^24^. In this paper, we provide a thorough evaluation of RDN and Soq in the context of Jurkat signalling and activation. By performing inhibitor treatments across three distinct models of T cell activation (soluble antibody crosslinking, APC-pMHC/TCR co-culture, and CD19-CAR/Raji co-culture), we find that RDN’s efficacy in abating T or CAR T cell signalling is higher than its efficacy in disrupting T cell activation markers. Further, Soq appears to thoroughly disrupt PLCγ1^Y783^ phosphorylation and IL-2 production even at low concentrations, while leaving CD69 expression intact. Finally, we find that while soluble antibody stimulation induces the most pY induction of any stimulation method, it is unable to activate T cells to the same degree as APC-pMHC/TCR or CD19-CAR/Raji stimulation. Thus, we provide the first phosphoproteomic analysis of RDN and Soq in the context of T and CAR T cell signalling and activation, thereby providing a molecular basis for their *in vivo* effects.

## 2 Results

### 2.1 Soq inhibits PLCγ1 phosphorylation without large-scale disturbance of the TCR signalling pathway

Due to the novelty and potential therapeutic uses of RDN and Soq, we aimed to thoroughly characterise their effects on Jurkat T and CAR T cell signalling and activation (Figure 1A). Isogenic Jurkat lines enable the replication of large-scale, deep phosphoproteomic analysis, thereby eliminating donor variability. We performed a titration of RDN and Soq on Jurkat T cells and assessed phosphorylation of key TCR signalling proteins after T cell activation by soluble antibody stimulation of the TCR and CD28, APC-pMHC/TCR co-culture, and CD19- CD28-ζ CAR/Raji (CD19-CAR/Raji) co-culture. We demonstrated that low concentrations (100 nM) of RDN or Soq were sufficient to reduce basal Erk1/2 phosphorylation (see Supporting Figure 1). We found that only 10 μM RDN was sufficient to impair phosphorylation of LAT^Y220^, a direct substrate of Zap70 ^25^ (Figure 1B-D). Phosphorylation of Lck^Y394^ was unaffected by RDN (Figure 1B, Supporting Figure 2), but the downstream pathway targets Erk1^T202Y204^/Erk2^T185Y187^ were significantly reduced at 1 and 10 μM RDN after 2.5 minutes of stimulation. During soluble antibody stimulation, Soq treatment did not significantly disrupt PLCγ1^Y783^ phosphorylation, a direct substrate of Itk, after 2.5 minutes. It minimally affected Erk1^T202Y204^/Erk2^T185Y187^ until a concentration of 10 μM Soq and did not affect LAT^Y220^ or Lck^Y394^ (Figure 1B-D, Supporting Figure 2). These trends remained true irrespective of the method of stimulation (Figure 1E-J, Supporting Figures 3-4), except for PLCγ1^Y783^, which was significantly impaired after Soq treatment in APC-pMHC/TCR and CD19-CAR/Raji conditions. Lck^Y394^ was not induced during APC-pMHC/TCR or CD19- CAR/Raji co-cultures (Supporting Figures 3 & 4). We also observed that phosphorylation of early TCR signalling proteins peaked around 2.5 minutes after stimulation, with a cool-down after 10 minutes (Figure 1, Supporting Figures 2-4, Supporting Tables 1-12). These data suggest that 10 μM RDN is required to disrupt TCR/CAR signalling, whereas 10 μM Soq or higher is required to achieve maximal Itk inhibition.

**Figure 1:**
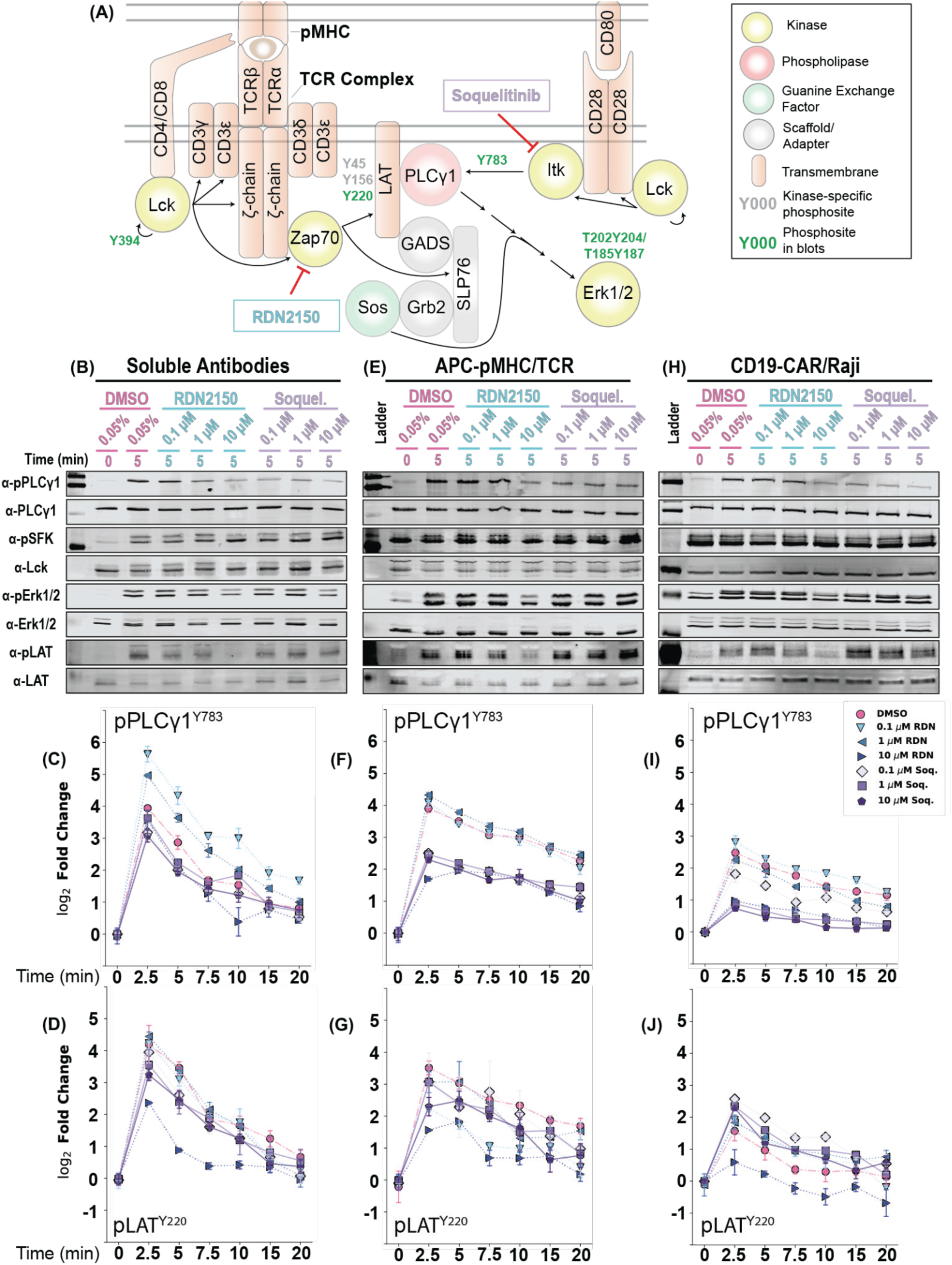
Soq inhibits PLCγ1^Y783^ while RDN inhibits both PLCγ1^Y783^ and LAT^Y220^ phosphorylation. (A) Schematic representation of T cell signalling, indicating the expected effects of inhibiting Zap70 (RDN) or Itk (Soq). Phosphorylation sites depicted in green are probed in subsequent Western blots. (B) Representative Western blots targeting PLCγ1^Y783^/PLCγ1, SFK^Y416^/Lck, Erk1^T202Y204^/Erk1 and Erk2^T185Y187^/Erk2, and LAT^Y220^/LAT in soluble antibody stimulation of the TCR (C) Quantification of PLCγ1^Y783^ during a time course soluble antibody stimulation of the TCR. (D) Quantification of LAT^Y220^ during a time course soluble antibody stimulation of the TCR. (E-G) As in (B-D), except for APC-pMHC stimulation of the Jurkat^OT-I^ TCR. (H-J) As in (B-D), except for Raji co-culture stimulation of a second-generation CD19-CAR. Statistical analysis for Western blots was performed using the Holm-Sidak correction of Fisher’s LSD pairwise comparisons and is available in Supporting Tables 1, 2, 5, 6, 9, and 10.

### 2.2 Both RDN and Soq inhibit IL-2 expression, but only RDN blocks CD69 upregulation

Signalling from the TCR/CAR is a rapid process, leading to the expression of new genes and the presentation/release of new proteins that elicit intercellular effects, such as cell death or the activation of neighbouring immune cells. T cell activation is a finely tuned process that requires precise, time-dependent activation of each component of the signalling pathway, and small perturbations can lead to over- or under-activation of T cells, ultimately resulting in autoimmunity^26^. To determine whether RDN or Soq impacted overall T cell activation, we evaluated expression of CD69 and IL-2, two activation markers of T and CAR T cells, after 24 hours of antibody-, pMHC-, and CD19-CAR/Raji-induced T cell activation. We found that soluble antibody stimulation mildly induced CD69 expression but failed to induce IL-2 expression (Figure 2A-B, Supporting Tables 13-14). Treatment with either RDN or Soq significantly reduced soluble antibody stimulation-induced CD69 expression at all tested concentrations (Supporting Table 13), but only 10 μM RDN completely ablated CD69 induction (Figure 2A). After APC-pMHC/TCR stimulation, CD69 and IL-2 were highly upregulated and significantly reduced after treatment with RDN (Figures 2C-D). CD69 expression was not affected by any tested concentration of Soq, but Soq treatment significantly reduced IL-2 production after APC-pMHC/TCR stimulation (Figures 2C-D, Supporting Tables 15-16). These trends were most evident in CD19-CAR/Raji stimulation conditions, as CD69 was widely expressed after co-culture and was reduced only by RDN treatment. IL-2 production was highest in CAR T cells (∼1000 pg/mL in DMSO control samples) and was significantly reduced with both RDN and Soq treatment (Figures 2E-F, Supporting Tables 17-18). These data in the context of the known TCR pathway ordering confirm that soluble antibody stimulation is insufficient to induce full T cell activation and demonstrate that the Itk-mediated branch of TCR signalling is responsible for IL-2 production.

**Figure 2:**
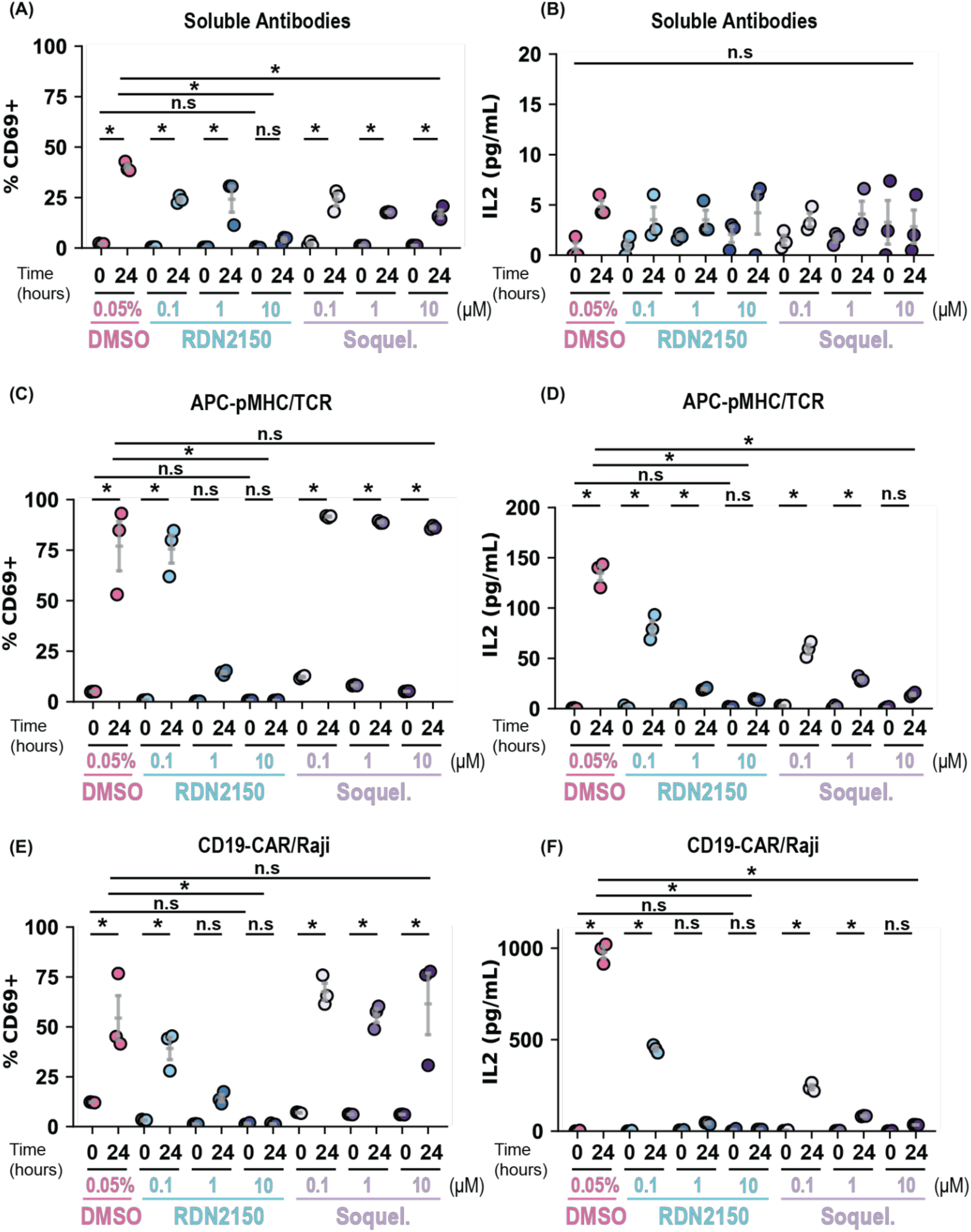
Interleukin-2 secretion, but not CD69 expression, is significantly reduced by Soq treatment. (A) Dotplot showing the percentage of CD69-positive cells measured by flow cytometry after 24 hours of soluble antibody stimulation. (B) Dotplot showing the concentration of IL-2 in the cell culture supernatant measured by ELISA after 24 hours of soluble antibody stimulation. (C-D) As in (A-B), except for APC-pMHC stimulation. (E-F) As in (A-B), except for Raji-co-culture stimulation of a second-generation CD19-CAR. Statistical analysis for flow cytometry and ELISA data was performed using a one-way ANOVA followed by the Holm-Sidak correction of Fisher’s LSD pairwise comparisons, if applicable, and is available in Supporting Tables 13-18.

### 2.3 RDN treatment reduces ligand-induced pY signalling more than Soq

To gain precise information regarding the roles of Zap70 and Itk in TCR- and CAR-mediated T cell activation, we leveraged the sensitivity of LC-MS/MS coupled with pY enrichment using the Src SH2 superbinder (sSH2) as previously described^27–31^. We sequenced a total of 773 T cell-specific pY sites in antibody-stimulated samples with high reproducibility (R>0.8 by multiple regression; see https://github.com/Aurdeegz/Zap70-Itk-Inhibitor-Profiling). Phosphotyrosine signalling from the TCR/CAR is well characterised, and Zap70/Itk have canonical roles in the progression of T cell activation (Figure 1A)^1,9,10,30,32–34^. Using our pY proteomics data, we observed that 2.5 minutes of soluble antibody stimulation of the TCR induced large-scale pY induction irrespective of RDN or Soq treatment (Figure 3A, top row, Supporting Figure 5). Known components of the TCR signalling cascade, including ITAMs on CD3ε and TCRζ, LAT^Y220^, Erk2^T185Y187^, Nck^Y105^, PLCγ1^Y775^, PLCγ1^Y783^, SFK^Y416,^ and Zap70^Y492^ were increased after 2.5 minutes with or without RDN and Soq. After 10 minutes, some of these pY sites returned to basal, although some pY sites, including ITAMs, Erk1^T202Y204^/Erk2^T185Y187^, PLCγ1^Y775^, and PLCγ1^Y783,^ remained elevated in DMSO and/or RDN treatment conditions, in line with our titration data (Figures 1, 3B, Supporting Table 19). Analysis of time course data using post-translational modification signature enrichment analysis (PTM-SEA), a GSEA-like algorithm for PTM proteomics data^35^, showed that soluble antibody stimulation strongly engaged TCR signalling kinases (Supporting Figure 6A), components of various signalling pathways (Supporting Figure 6B), and Perturbation Signatures associated with cell activation (Supporting Figure 6C). In contrast, APC-pMHC/TCR stimulation modestly induced pY signalling by Western blot and proteomics for pY, pLAT, and pPLCγ1 (Figure 3A, middle row, Supporting Figure 7). The effects of RDN treatment were more evident with fewer significantly induced pY sites after 2.5 minutes (365 total pY sites, Figure 3A, middle row). APC-pMHC/TCR stimulation did not uniformly induce phosphorylation along CD3/TCRζ ITAMs, and Soq treatment did very little to reduce pY induction along the T cell signalling pathway (Figure 3C, Supporting Table 20). In agreement with the antibody stimulation data, APC-pMHC/TCR stimulation significantly upregulated Kinase Signatures (Supporting Figure 8A), Pathway Signatures with the notable exception of TCR signalling (Supporting Figure 8B), and perturbation signatures associated with cell activation, including ‘ANTI CD3’ and ‘IL 2’ (Supporting Figure 8C). Interestingly, CD19-CAR/Raji co-culture significantly increased few pY sites (299 total pY sites, Figure 3A, Supporting Figure 9) after 2.5 minutes with TCR signalling pY sites like Erk2^T185Y187^, Nck1^Y105^, PLCγ1^Y775^, PLCγ1^Y783^, and Zap70^Y492^ being among them in the DMSO control (Figure 3D, Supporting Table 21). Treatment with RDN ablated the induction of all TCR signalling pY sites at 2.5 minutes after co-culture. In contrast, Soq treatment had little effect on pY induction along the TCR signalling pathway (Figure 3D). Finally, although many Kinase Signatures (Supporting Figure 10A), Pathway Signatures (Supporting Figure 10B), and Perturbation Signatures (Supporting Figure 10C) were significantly upregulated in CD19-CAR/Raji stimulations with DMSO or Soq treatment, almost no signatures were significantly induced during RDN treatment, supporting that Zap70 is integral to the CD19-CAR activation^36,37^. These data, in combination with Figure 1, support the notion that pY induction is heavily dependent on the form of stimulation, and that the effectiveness of RDN and Soq treatment on pY signalling varies greatly with stimulation strength.

**Figure 3:**
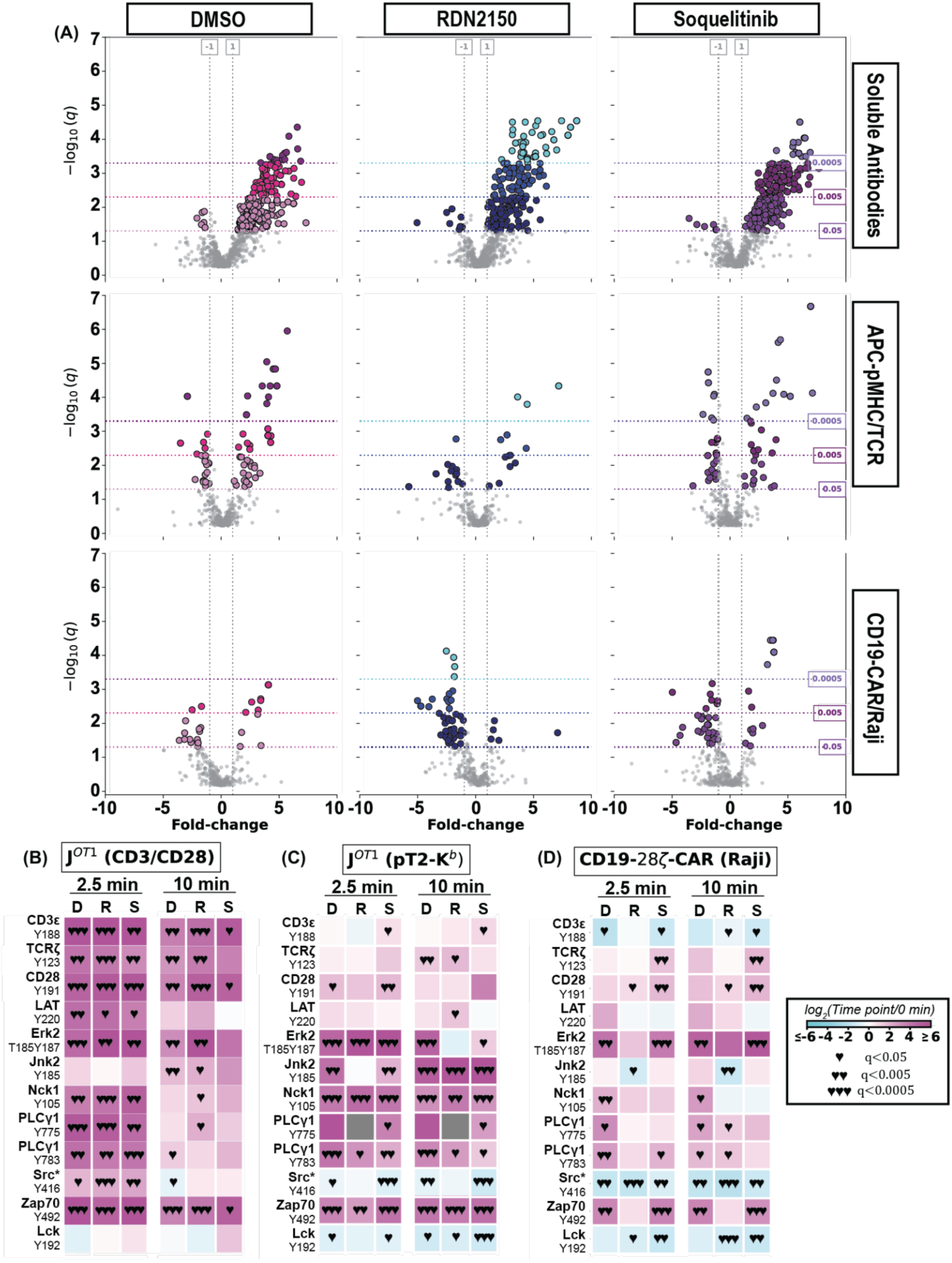
Antibody/TCR stimulation massively induces tyrosine phosphorylation irrespective of RDN or Soq treatment. (A) Volcano plot analysis of pY proteomics data comparing 2.5 minutes versus 0 minutes of antibody stimulation (top), APC-pMHC/TCR stimulation (middle), and CD19-CAR/Raji (bottom) in DMSO (left), RDN (middle), and Soq (right) treatment conditions. (B-D) Select site-specific heatmaps of time course pY proteomics data after antibody stimulation, APC-pMHC/TCR stimulation, and CD19-CAR/Raji stimulation. ♥ indicates q < 0.05, and ♥♥ indicates q < 0.005, and ♥♥♥ indicates q < 0.0005.

### 2.4 RDN treatment disrupts the TCR pathway more than Soq treatment

Given the ability of cells to retain signalling capacity in the presence of each inhibitor, we wished to evaluate how inhibition affected pY-site abundance relative to the DMSO control. As expected, treatment with either 10 μM RDN or 10 μM Soq uniformly reduced pY abundance in soluble antibody-treated samples, with the majority of pY sites being significantly reduced at 0, 2.5, and 10 minutes of stimulation when compared to the DMSO control (Figure 4A, Supporting Figure 5, Supporting Table 19). pY sites along the TCR signalling pathway, including the Erk1/2 activation sites, Jnk2/3 activation sites, and multiple sites on PLCγ1 and Zap70 were reduced during RDN treatment. Interestingly, 10 μM RDN treatment elevated Zap70^Y493^ abundance at the basal and 10-minute stimulation time points, while reducing various TCR ITAM sites, such as TCRζ^Y64^/TCRζ^Y72^ and CD3ε^Y188^ (Figure 4B, Supporting Table 19). PTM-SEA showed massive downregulation of activating perturbation signatures like ‘ANTI CD3’ and ‘IL 2’ (Figure 4C), and upregulation of inhibitor signatures like ‘ERLOTINIB’ and ‘U0126’ (Figure 4D), suggesting that the phosphoproteomic signature of RDN and Soq mimicked that of pre-established inhibitors. Our results for RDN treatment in APC-pMHC/TCR and CD19-CAR/Raji stimulations were similar, showing that the majority of significantly changing pY sites were reduced (Supporting Figures 11A, 12A), those sites often were located on TCR signalling pathway proteins (Supporting Figures 11B, 12B), and that the PhosphositePlus Perturbation signatures and NetPath Pathway signatures were, in general, significantly downregulated during inhibitor treatment (Supporting Figures 11C-D, 12C-D). In contrast, pY site abundance changes during Soq treatment were more evenly distributed in APC-pMHC/TCR and CD19- CAR/Raji co-cultures, with very few TCR signalling proteins showing reduced pY site abundance (Supporting Figures 11, 12). Importantly, we consistently observed PLCγ1^Y783^ as significantly reduced compared to the DMSO control in APC-pMHC/TCR and CD19- CAR/Raji co-cultures (Supporting Figures 7, 9, 11, 12), indicating that Soq and RDN were present and active. Together, these data suggest that RDN, more than Soq, reduces pY abundance in T cells across stimulation methods.

**Figure 4:**
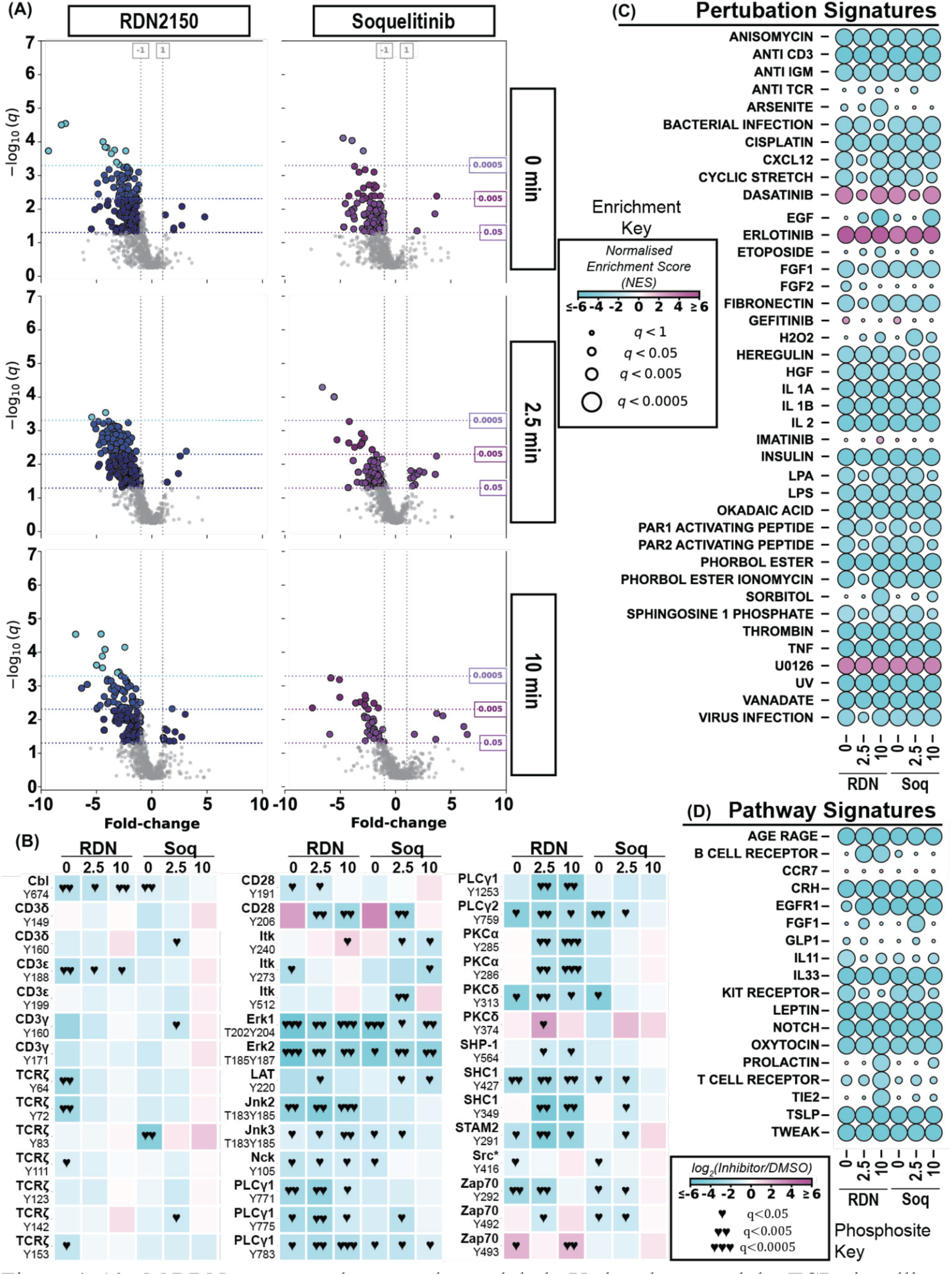
10 uM RDN treatment downregulates global pY abundance and the TCR signalling pathway more than Soq treatment. (A) Volcano plot analysis of pY proteomics data comparing RDN and Soq treatment to a DMSO control in samples stimulated with soluble antibodies for 0 (top row), 2.5 (middle row), and 10 (bottom row) minutes. (B) Heat maps comparing RDN and Soq treatment to a DMSO control for specific pY sites in the T cell signalling pathway. ♥ indicates q < 0.05, and ♥♥ indicates q < 0.005, and ♥♥♥ indicates q < 0.0005. (C-D) PTM-SEA analysis of phosphoproteomic data showing PhosphoSitePlus Perturbation Signatures and NetPath Pathway Signatures, respectively.

### 2.5 Stimulation method-dependent effects on pY abundance during RDN and Soq treatment

Considering that RDN and Soq both significantly reduced pY abundance on T cell signalling proteins (Figure 4, Supporting Figures 11, 12), yet had different effects on CD69 expression and IL-2 secretion (Figure 2), we sought to characterise the broader impact of RDN and Soq treatment. Treatment with RDN led to the greatest number of significantly differentially expressed pY sites (q<0.05 compared to DMSO control) at all time points in our soluble antibody stimulation model (219 vs 118, 277 vs 137, 174 vs 48 for 0-minute, 2.5-minute, and 10-minute, respectively). Most of those sites were unique to RDN treatment (Figure 5A-C). This trend held for APC-pMHC/TCR (Figures 5D-F) and CD19-CAR/Raji (Figures 5G-H) stimulations, particularly post-stimulation. Fewer than 50% of differential pY sites identified in RDN treatment were shared with Soq treatment (except at 0 minutes of antibody stimulation; Figures 5A-I). These data suggest that, despite similar trends of pY abundance in TCR signalling proteins compared with the DMSO control (Figure 4, Supporting Figures 11, 12), RDN and Soq control unique subsets of pY sites in Jurkat T cells. In soluble antibody stimulation samples, we observed differential, non-canonical pY sites on key TCR signalling proteins, which were significantly reduced in both RDN and Soq, such as Lck^Y470^ and Fyb^Y803^. Some TCR pY sites were significantly reduced only in RDN-treated samples, such as Cofilin-1^Y85^, PAG^Y317^, PAG^Y387^, SHP-1^Y536^, and Zap70^Y164^, whereas others, including Itk^Y588,^ were significantly reduced during Soq treatment. Notably, we observed that pY sites on heterogeneous nuclear ribonucleoproteins (hnRNP) F^Y82^, hnRNP K^Y280^, and hnRNP H1^Y82^ were significantly reduced during RDN treatment. The hnRNP family of proteins is involved in stabilising pre-mRNA during mRNA maturation, and thus directly influences protein translation^38^. While the function of these pY sites is unknown, hnRNP F^Y82^ flanks a quasi-RNA recognition motif (RRM), hnRNP H1^Y82^ lays inside a qRRM, and hnRNP K^Y280^ is within an arginine-rich patch, which may be involved in phase separation of hnRNPs^38–42^.

**Figure 5:**
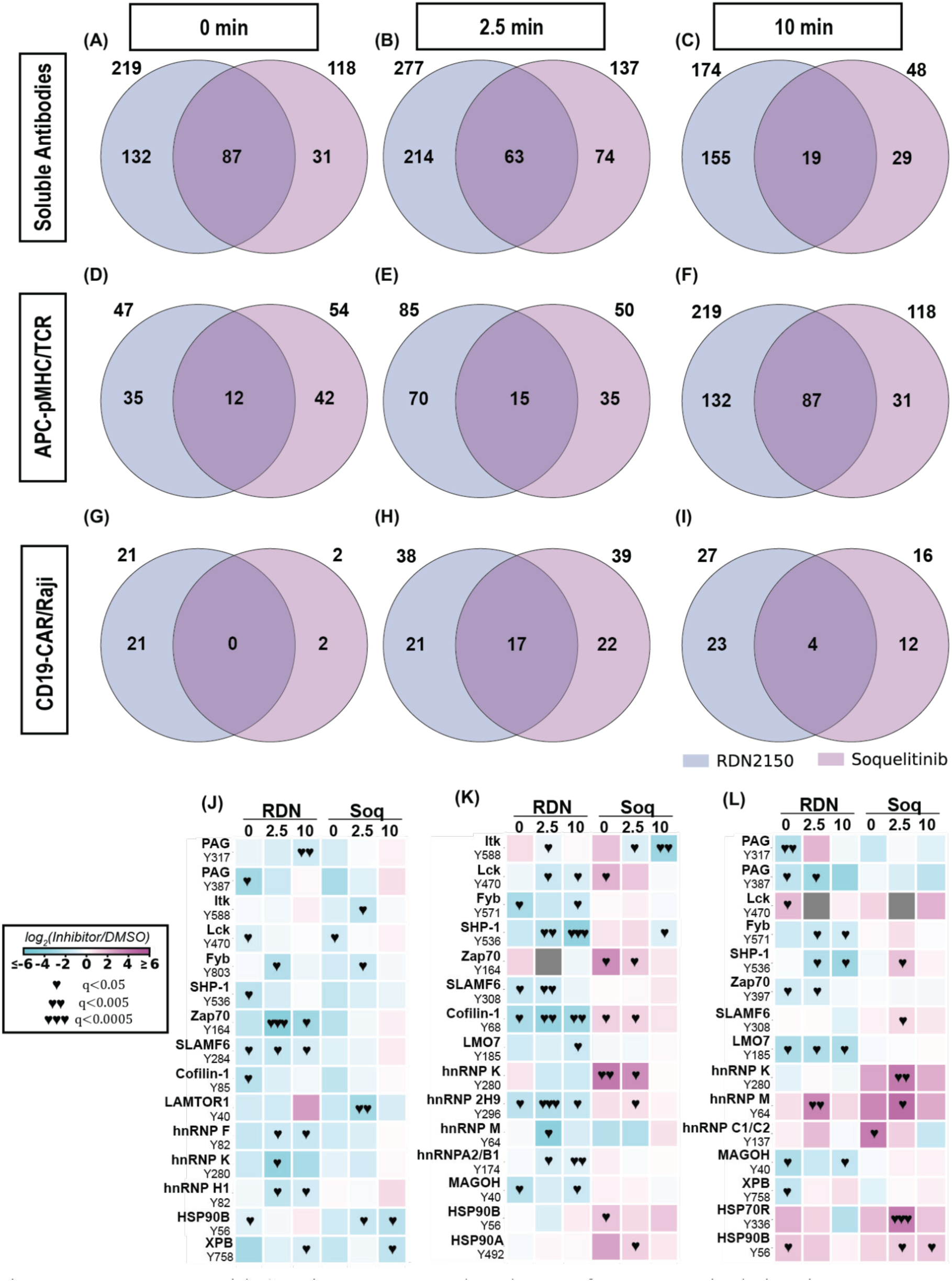
Treatment with Soq increases pY abundance of non-canonical sites in response to APC-pMHC/TCR and CD19-CAR/Raji stimulation. (A-C) Venn diagrams showing overlap of unique pY sites observed as significantly changing (q < 0.05) compared to DMSO for soluble antibody stimulated samples at 0 minutes, 2.5 minutes, and 10 minutes, respectively. (D-F) As in A-C, except for APC-pMHC/TCR stimulated samples. (G-I) As in A-C, except for CD19-CAR/Raji stimulated samples. (J-L) Heat maps comparing RDN and Soq treatment to a DMSO control for specific, non-canonical pY sites in antibody, APC-pMHC/TCR, and CD19-CAR/Raji stimulation, respectively. ♥ indicates q < 0.05, and ♥♥ indicates q < 0.005, and ♥♥♥ indicates q < 0.0005.

Similarly, many non-canonical pY sites on TCR signalling proteins were reduced more in RDN treatment than Soq treatment during APC-pMHC/TCR stimulation (Figure 5K), and some pY sites were even increased in Soq treatment compared to the DMSO control, including Lck^Y470^, Zap70^Y164^, and Cofilin-1^Y68^. Notably, the pY sites hnRNP 2H9^Y296^, hnRNP M^Y64^, and hnRNP A2/B1^Y174^ were significantly reduced by RDN treatment in APC-pMHC/TCR treatment. However, hnRNP K^Y280^ and hnRNP 2H9^Y296^ were significantly increased during Soq treatment (Figure 5K). While the trends for RDN treatment reducing non-canonical pY sites on signalling proteins held for CD19-CAR/Raji co-culture, we observed an elevation of hnRNP M^Y64^ in RDN treatment and hnRNP K^Y280^ (2.5 min), hnRNP M^Y64^ (2.5 min), and hnRNP C1/C2^Y137^ (0 min) during Soq treatment (Figure 5L). Altogether, our data demonstrate that Zap70 is the major regulator of main-line T cell signalling; however, Zap70 and Itk control unique, stimulation-dependent branches of the non-canonical T cell pY proteome.

## 3 Discussion

Activation via the T cell receptor and, more recently, chimeric antigen receptors has been extensively studied in Jurkat cells and primary T cells. While Jurkat T cells are known to be deficient in PTEN, resulting in constitutive Itk localisation to the membrane^43^, researchers use Jurkat T cells due to their isogenic nature, which reduces donor/organismal variability and allows for greater pathway insight. Using Jurkats and primary T cells, researchers demonstrate that the basic TCR/CAR pathway involves Lck/Fyn-mediated phosphorylation of ITAMs on CD3 and/or TCRζ, Zap70 phosphorylation, and activation. Subsequently, the production of a signalosome, including LAT/SLP76 in traditional T cells or CAR/SLP76 in CAR T cells, leads to signal diversification. However, while T and CAR T cell signalling is well studied^1,5^, many integral aspects of TCR/CAR signalling remain elusive. For example, researchers have only recently decoded the role of Zap70 in the kinetic gatekeeping of T cell activation, how Lck and Zap70 coordinate to promote LAT phosphorylation, and how the Itk substrate site GADS^Y45^ integrates CD28 costimulation with TCR stimulation. Similarly, recent construction of optimized CARs that include unintuitive intracellular domains identified through high-throughput screens in Jurkat cells^17–22^ and point mutations in CARs that can improve function^9,10,12^ are the first steps toward bringing this promising immunotherapy to more patients. Notably, signalling studies of CARs and TCRs are often confounded by the choice of stimulation method (e.g., antibodies, reconstituted MHCs, cell-to-cell contact), making results difficult to compare and impossible to determine stimulation- method-dependent effects on activation. Together, while we have made significant advances in understanding TCR and CAR signalling, our basic understanding of the processes underlying T cell activation through these receptors can be improved.

Here, we aimed to leverage the sensitivity of mass spectrometry-based phosphotyrosine proteomics to determine the role that Zap70 and Itk kinase activity play in Jurkat T and CAR T cell activation by using two novel T cell inhibitors. The first, RDN, was described by Rao *et al*. (2023) and shown to covalently modify Zap70^C346^, which lies at the N-terminus of the catalytic domain of Zap70. Rao *et al*. showed that RDN suppressed *in vitro* kinase activity more than 98% at concentrations of 1 μM, suppressed LAT^Y220^ phosphorylation in Jurkats at concentrations as low as 400 nM with 48 hours of CD3 stimulation (albeit without replicates), and suppressed CD69, CD25, IFN-γ, and IL-17A production at concentrations between 100-400 nM in primary murine CD4+ T cells^23^. While our functional activation data are in agreement, showing that 100-1000 nM is effective for suppressing CD69 and IL-2 production (Figure 2), we found no significant decrease in LAT^Y220^ phosphorylation until 10 μM RDN after less than 20 minutes of CD3/CD28 crosslinking (Figure 1). Further, we found that treatment with low concentrations of RDN could significantly improve PLCγ1^Y783^ induction after soluble antibody stimulation (Figure 1C) and Erk1^T202Y204^/Erk2^T185Y187^ induction after any stimulation method (Figure 1B, Supporting Figures 2-4). These data are consistent with previous findings showing that the kinase domain of Zap70 mediates basal signalling and negative feedback on the TCR to ensure proper T cell signalling and activation. With a treatment of 10 μM RDN, our pY proteomics data showed that the majority of pY sites were significantly decreased when compared to a DMSO control, many of which are on integral T cell proteins like Cbl, CD28, Erk1, Erk2, LAT, Jnk2, Jnk3, Nck, PLCγ1, PKCα, PKCδ, SHP-1, SHC1, and STAM2. Further, the molecular signatures in the data resembled that of other kinase inhibitors and showed significant downregulation of signatures associated with cell signalling pathways (Figure 4). Despite this, soluble antibody stimulation was able to induce wide-scale pY in the presence of 10 μM RDN. In contrast, stimulation with APC-pMHC/TCR or CD19-CAR/Raji co-culture appeared muted in the presence or absence of RDN (Figure 3). The residual pathway stimulation in the presence of either Soq or RDN (Figure 3) could reflect partial inhibition of these kinases at the concentrations tested or compensatory pathways through parallel kinases such as Lck or Fyn. Together, these data in Jurkat cells support a model by which Zap70 is a primary driver of pY signalling from the TCR, is more effective in systems with weaker pY induction, and inhibits the production of activation markers even when insufficient to block pY signalling completely (Figure 6), in line with the current model where Zap70 acts as a kinetic proofreader of TCR signalling.

**Figure 6:**
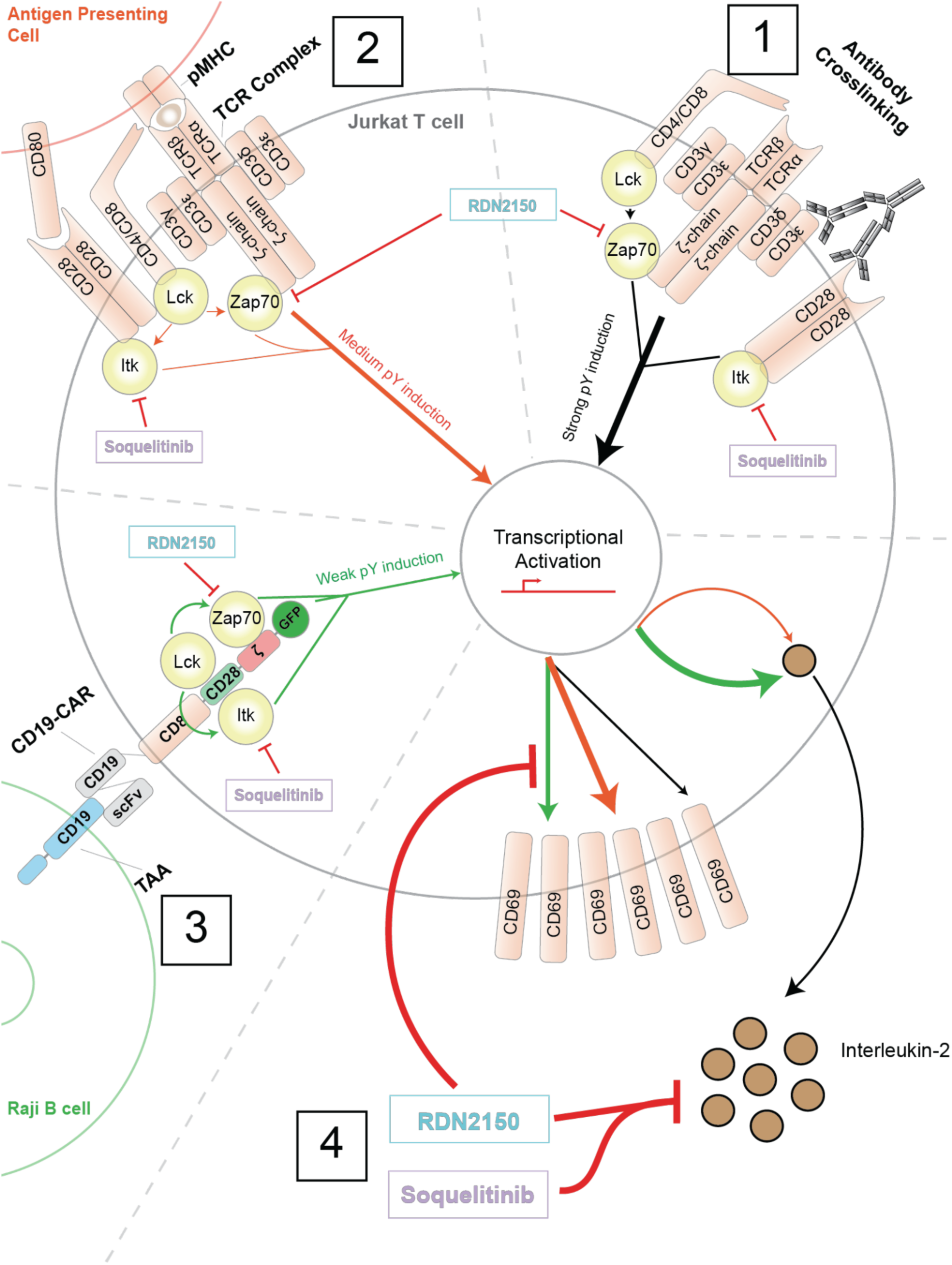
pY induction and the T cell activation markers CD69/IL-2 are inversely related. [Part 1] Soluble antibody crosslinking of CD3ε and CD28 induces very strong pY signalling down the T cell signalling cascade. Treatment with RDN or Soq reduces pY abundance at 0, 2.5, and 10 minutes post-stimulation (Figures 1, 3, & 4); however, pY signalling remains induced. [Part 2] CD19-CAR/Raji binding through co-culture induces TCR signalling to a milder degree than antibody crosslinking, and RDN largely impairs pY induction (Figure 1, 4). [Part 3] CD19-CAR/Raji binding induces much weaker pY signalling than either antibody crosslinking or APC-pMHC/TCR binding. Treatment with RDN blunts pY signalling entirely (Figures 1, 4). [Part 4] APC-pMHC/TCR and CD19-CAR/Raji co-culture induce high CD69 expression, whereas soluble antibody crosslinking does not, and soluble antibody crosslinking is unable to induce IL-2 production (Figures 2A, 2C, 2E). RDN, but not Soq, blocks CD69 expression; however, both RDN and Soq block IL-2 production (Figure 2).

Next, we evaluated the Itk inhibitor Soquelitinib (Soq; formerly CPI-818), developed by Corvus Pharmaceuticals and in clinical trials for T cell lymphoma treatment^48^. Recently, Hsu *et al*. (2024) described the synthesis and characterisation of Soq, which covalently modifies Itk^C442^, a residue at the heart of the Itk kinase domain. Hyu *et al*. showed that Itk prevented CD3ε-induced PLCγ1^Y783^ phosphorylation at concentrations as low as 100 nM, impaired IL-2 production at 0.1-1 μM, significantly decreased primary cell viability at concentrations greater than 1 μM, and skewed T cell differentiation towards a Th1 phenotype. Our data largely agreed with Hsu *et al*., showing that low concentrations of Soq impaired PLCγ1^Y783^ and Erk1^T202Y204^/Erk2^T185Y187^ phosphorylation in response to CD3ε/CD28, TCR, and CAR stimulation (Figure 1, Supporting Figures 2-4), as well as IL-2 production (Figure 2).

Interestingly, we found that with APC-pMHC/TCR and CD19-CAR/Raji stimulation, all tested concentrations of Soq were insufficient to reduce activation-induced CD69 expression, whereas under soluble-antibody stimulation, it did significantly reduce CD69 upregulation. (Figure 2). Previous literature has shown that the kinase activity is important for IL-2 production, although the mechanism by which Itk controls IL-2 production is unclear^49^, and that CD69 expression is an ‘all or nothing’ threshold that depends on ligand strength^50^. Thus, while our data in Jurkat cells showed that Itk is involved in CD69 production, Itk was less critical than Zap70 kinase activity, which was necessary for both CD69 and IL-2 production (Figure 2). Using a treatment of 10 μM Soq, we found that global pY abundance decreased compared to a DMSO control; however, pY sites along the T cell signalling pathway did not appear to be heavily affected. Notably, PLCγ1^Y783^ and Erk1^T202Y204^/Erk2^T185Y187^ were reduced, in line with our titration Western blot data, and some pY sites on Zap70 were significantly reduced. The molecular signatures associated with Soq treatment reflected its inhibitory activity (Figure 4). In the context of pY induction, Soq appeared to have little effect, regardless of the stimulation method (Figure 3A). In total, our data support a model of Itk function, whereby Itk acts as a rheostat to tune T cell activation, with particular control over the IL-2 production pathway (Figure 6).

Lastly, our data provide a comprehensive comparison of T cell activation methods frequently used in the literature, including soluble antibody stimulation of the TCR, APC-pMHC/TCR co-culture, and CD19-CAR/Raji co-culture. We found that fold induction of key TCR signalling pY sites by Western blot differed between soluble antibody, APC-pMHC/TCR, and CD19-CAR/Raji stimulations (Figure 1), and this difference did not correlate with CD69 expression or IL-2 secretion (Figure 2). We found that soluble antibody stimulation acted like a molecular sledgehammer on the pY proteome (Figures 1 and 3); however, it was unable to sufficiently promote CD69 expression or IL-2 secretion compared to APC-pMHC/TCR or CD19-CAR/Raji co-culture. While RDN and Soq reduced pY abundance on T cell signalling proteins irrespective of stimulation method (Figure 4), they had unique effects on non-canonical TCR pY sites. In particular, RDN treatment reduced phosphorylation of multiple heterogeneous nuclear ribonucleoproteins (hnRNPs) independent of stimulation method; however, Soq treatment increased pY abundance on multiple hnRNP pY sites in APC-pMHC/TCR and CD19-CAR/Raji co-culture experiments. While many of these pY sites are uncharacterised, hnRNPs play a significant role in T cell physiology. In particular, Chang *et al*. (2009) showed that hnRNP K was a direct pathway target of the TCR, was required for effective IL-2 production, and promoted Vav1 proteolysis^51^. Meininger *et al*. (2016) showed that hnRNP U suppressed exon 7 inclusion in *MALT1* mRNA splicing, thereby regulating MALT1A expression and T cell activation^52^. White *et al*. (2025) showed that hnRNP L controlled differentiation of peripheral T cells, particularly into T follicular helper cells^53^.

Finally, Shankarling *et al*. (2014) used crosslinking and immunoprecipitation sequencing to identify alternative splicing events of TCR signalling mRNAs in response to TCR signalling^54^. Although the signalling pathway linking T cell signalling and hnRNP regulation remains unclear in many cases, our data suggest that Zap70 and Itk play stimulation-dependent roles on the abundance of hnRNP pY sites.

Here, we provide a thorough, site-specific resource on the activation pathway of Jurkat T and CAR T cells in the presence of two T cell inhibitors, RDN and Soq. We showed that Zap70 is the dominant regulator of TCR signal initiation and T cell activation, while Itk selectively regulates IL-2 induction through a more limited branch of T cell signalling. While our studies directly inform the interpretation of high-throughput Jurkat CAR engineering screens, future targeted studies with primary human T cells will define the translational potential of these findings to improve CAR T therapies and identify the possible benefits of combination therapies.

## 4 Materials and Methods

### 4.1 Cell culture and SILAC labelling

Jurkat T cells (clone E6.1), J^OT1^ (Jurkats expressing murine OT1 TCR and human CD8; gift from Dr. Arthur Weiss), 28ζ-CAR (Jurkats expressing a FMC63 scFv, CD8 hinge/transmembrane, CD28 costimulatory domain, TCRζ stimulation domain containing CAR. Characterised in Callahan *et al*. (2025)), T2-K^b^ cells (T2 cells expressing the Kb class 1 MHC; gift from Dr. Arthur Weiss), and Raji B cells (gift from Dr. Xiaolei Su), were maintained in RPMI 1640 supplemented with 10% FBS (Peak Serum #PS-FB3 Lot 01Z2233), 1X penicillin streptomycin glutamine (Cytiva #SV30082.01), and 2.5 μg/mL plasmocin (InvivoGen #ant-mpp) in a humidified incubator at 37 C, 5% CO_2_. For co-culture phosphoproteomics, Jurkat T cell derivatives were grown initially in supplemented RPMI 1640 as described above, before transitioning to SILAC RPMI (Thermo #A33823) supplemented with 10% dialysed FBS, 1X penicillin streptomycin glutamine, and 2.5 μg/mL plasmocin, 0.38 mM ^13^C_6_ ^15^N_4_ Arginine (Millipore #608033), and 0.22 mM ^13^C_6_ ^15^N_2_ Lysine (Cambridge Isotopes #CNLM-291-H-1) for at least 7 days, as previously described^30,31^.

### 4.2 Inhibitor treatment

For inhibitor treatment experiments, Jurkat T cells were collected by centrifugation at 400 xg for 5 minutes, washed once in unsupplemented RPMI 1640 (or SILAC RPMI 1640 for phosphoproteomics), then resuspended at a concentration of 1x10^7^ cells/mL in RPMI 1640 supplemented with the indicated concentration of RDN (MedChemExpress #HY-155978), Soq (MedChemExpress #HY-150298), or DMSO and were allowed to incubate at 37 C, 5% CO2 in a humidified incubator for 3 hours. After inhibitor treatment, the cells were collected and washed using 1X DPBS supplemented with inhibitor, then resuspended at a concentration of 2x10^8^ cells/mL for stimulation.

### 4.3 T cell receptor/Chimeric Antigen Receptor stimulation

Soluble antibody stimulation of the T cell receptor was performed as previously described^29^. Briefly, Jurkat T cells were suspended in RPMI 1640 at a concentration of 2x10^8^ cells/mL, then allowed to rest at 37 C for 30 minutes. T cell receptor stimulation was initiated by addition of α-CD3ε (Thermo #16-0037-85, clone OKT3; 2 μg/mL final concentration) and α- CD28 (Thermo #16-0288-81, clone CD28.6; 2 μg/mL final concentration) antibodies for 30 seconds, followed by addition of α-Mouse IgG (Jackson ImmunoResearch #115-005-062, 22 μg/mL final concentration). Stimulation was stopped by addition of 9 M Urea Lysis Buffer (9 M Urea, 1 mM sodium orthovanadate, 1 mM sodium pyrophosphate, 1 mM β- glycerophosphate in 20 mM HEPES) at the indicated time point.

Co-culture stimulation of Jurkats expressing the OT1 TCR was performed by loading T2-K^b^ cells with OVA^257-264^ (SIINFEKL; Rockland #000-001-M45) by incubation at plain RPMI 1640 supplemented with 1 μM OVA^257-264^ peptide for 3 hours, then washing in 1X DPBS and resuspending at 200,000,000 cells/mL. Stimulation was initiated by the 1:1 (v/v) addition of OVA^257-264^ loaded T2-K^b^ cells to JOT1 cells followed by a 30-second centrifugation at 500 xg to initiate cell-to-cell contact. Stimulation was stopped as described above.

Co-culture stimulation of 28ζ-CAR T cells was performed as previously described^28,31^ by 1:1 addition of Raji B cells to 28ζ-CAR T cells, centrifugation at 500 xg for 30 seconds, and lysis with 9 M Urea Lysis Buffer. Samples for Western blotting were then sonicated and diluted 1:1 in 2x Laemmli sample buffer, as previously described^29^.

### 4.4 Flow cytometry

T cell activation was assessed by surface expression of CD69 after TCR stimulation as previously described^28^. Briefly, cells were treated with the inhibitor as described in 2.2, except for one hour, then activated as described in 2.3 using soluble antibodies or PFA fixed target cells, then were allowed to incubate at 37 °C, 5% CO_2_ in a humidified incubator for 24 hours. After 24 hours, about 1x10^6^ T cells (2x10^6^ total) were collected by centrifugation at 400 xg for 5 minutes, then resuspended in flow cytometry buffer (2% FBS in 1X DPBS) and treated with APC-conjugated α-CD69 antibody (1:50; BD Pharmingen #560967) for 30 minutes at 4 C. After antibody treatment, cells were washed thrice in 1 mL flow cytometry buffer, fixed with 2% PFA for 15 minutes in the dark, then washed thrice in 1 mL flow cytometry buffer before analysis on a Cytek Aurora flow cytometer. Output files were analysed using Floreada.io, a free online tool for flow cytometry analysis. The cell culture supernatant from these experiments was saved and assessed for IL-2 by ELISA (section 2.6).

### 4.5 Western blotting

Western blotting was performed as previously described^28–31^. Briefly, lysates were separated on 10% polyacrylamide gels poured in house at 100 V for 120 minutes, then transferred to PVDF membranes (Millipore #IPFL00010) at 100 V for 100 minutes using Tris-Glycine running and transfer buffers. Membranes were blocked in Intercept TBS Blocking Buffer (LI-COR #927-60003) for 1 hour at room temperature then incubated with primary antibodies diluted in 5% BSA in 1x TBST overnight at 4C. Primary antibodies solution was removed, the membranes were washed three times with 1x TBST, then incubated with secondary antibodies diluted in 5% BSA in 1x TBST for 1 hour at room temperature. After secondary incubation, the membranes were washed three times with 1x TBST, once with 1x TBS, then imaged on a LI-COR OdysseyM imager. Quantification of Western blots was performed as previously described^30^.

### 4.6 Interleukin-2 ELISA

Samples for IL-2 ELISAs were taken from the cell culture supernatant during flow cytometry preparation and frozen at -80 C until use. Interleuken-2 abundance was assessed using the Human IL-2 Sandwich ELISA Kit (proteintech #KE00017) per the manufacturer’s instructions. Briefly, 100 uL of samples diluted 1:4 in complete growth media were incubated in the ELISA plate for 2 hours at room temperature and washed four times in Wash Buffer before addition of 100 uL Detection Antibody solution. The plate was incubated at room temperature for 1 hour with Detection Antibody Solution, then washed four times with Wash Buffer. Next, 100 uL of Streptavidin-HRP solution was added to each well and incubated at room temperature for 40 minutes before washing four times with Wash Buffer. For signal development, 100 uL of TMB was added to each well and incubated for 15 minutes before quenching with 100 uL of Stop Solution. Absorbance was read at 450 nm with correction at 630 nm. Absolute quantitation was determined by using a standard curve of Human IL-2.

### 4.7 Sample preparation and phosphoproteomic sample acquisition

#### 4.7.1 Reduction, alkylation, trypsinization and desalting

Eleven milligrams of total protein were taken for each sample, then samples were adjusted to an equivalent volume (4.45 mL, ∼1.6 M Urea final) with 20 mM HEPES, then reduced with 10 mM dithiothreitol (final) for 30 minutes at 37 C, then alkylated with 10 mM iodoacetamide (final) for 15 minutes in the dark. In solution trypsinisation was initiated with the addition of Sequencing Grade Trypsin (Promega #V5113) at a ratio of 1:100 trypsin:protein. After trypsinisation, samples were acidified to 1% trifluoroacetic acid (TFA), then cleared by centrifugation. Samples were desalted using Waters C18 vacuum columns as previously described^29^. Under low vacuum, columns were activated using 100% acetonitrile (ACN), then equilibrated with 2 washes of 0.1% TFA. Samples were then passed through the columns, trapping peptides, and washed three times with 1, 5, and 6 mL of 0.1% TFA, before elution with 40% TFA and 0.1% ACN. After elution, 10 pmol of a phosphotyrosine standard (LIEDpYTAK) was added to each sample, then each sample was diluted to 20% ACN before freezing and 48-hour lyophilisation.

#### 4.7.2 pY enrichment with Src SH2 superbinder

Phosphotyrosine enrichment using recombinant Src SH2 superbinder (sSH2) was performed as previously described^27–31^. Briefly, ∼11 mg of dried peptide was resuspended in ice-cold IAP buffer (10 mM sodium phosphate monobasic monohydrate, 50 mM sodium chloride, 50 mM morpholinopropanesulfonic acid) at room temperature for 30 minutes with brief, interspersed vortexing. One hundred micrograms of recombinant sSH2 conjugated to agarose beads in-house was aliquoted into microcentrifuge tubes, then washed 3 times (1,500 xg, 2 min to collect beads) with 1 mL of ice-cold IAP buffer before addition of sample for 2 hours at 4 °C on an end-over-end rotator. After pY peptide binding, beads were washed 2 times with 1 mL ice-cold IAP buffer, then 3 times with ice-cold HPLC water. After the last water wash, all of the liquid was removed from the beads using an insulin syringe. Phosphotyrosine peptides were eluted from the beads using 0.15% TFA on a thermomixer at 1,150 rpm for 10 minutes, twice. Eluted pY peptides were then desalted using Empore C18 StageTips as previously described^29^ and dried by speed-vac before LC-MS/MS acquisition.

#### 4.7.3 LC-MS/MS acquisition

Samples were separated as previously described^28,29^ on a Vanquish Neo uHPLC system with an Acclaim PepMap C18 Trap column (7 cm, 2 cm bed length, 3 μm particle size, 100 A) and IonOpticks Aurora Ultimate (25 cm long x 75 μm ID, 1.7 μm particle size) and acquired on an Orbitrap Ascend mass spectrometer (Thermo Fisher). The separation gradient started at 5% Solvent B (80% ACN, 0.1% Formic Acid [FA])/95% Solvent A (0.1% FA) and increased to 30% Solvent B over 50 minutes, then spiked to 99% Solvent B for the remaining 5 minutes of acquisition. After acquisition, the flow path was washed with 4 alternating zebra washes, alternating between 100% Solvent A and 100% Solvent B, before being reequilibrated for the next run. The spray voltage was 2050 V, with an ion transfer tube temperature of 275 °C, and a lock mass of 445.12003. For MS1 acquisition, wide quant isolation on the Orbitrap detector at a resolution of 120,000 with a mass range of 300-1500 m/z was used. The maximum injection time was 50 ms, the normalised AGC target was 250%, and the RF lens was set to 30%, and data collection was in centroid mode. The following filters were included: monoisotopic peak and peptide mode, Charge state (+2-+7), Precursor Selection Range (375- 1,500 m/z), Dynamic Exclusion (exclude after 1 time, 15 sec exclusion time, 10ppm low and high mass tolerance, excluding isotopes), Intensity Threshold (intensity range of 20000 (min) – 1E20 (max) with relative intensity threshold of 20%). MS2 spectra were acquired in data-dependent mode with a cycle time of 3 seconds, normalised HCD fragmentation energy of 30%, and a 0.7 m/s quadrupole isolation window. Peptide fragments were detected using the ion trap detector with Turbo Mode active (scan range 150-2000 m/z) in centroid mode with the following settings: 35 ms maximum injection time, 100% normalised AGC target, dynamic maximum injection time active.

#### 4.7.4 Data analysis, database searching and relative abundance quantification

.RAW files from the mass spectrometer were searched as previously described using the High Throughput Autonomous Proteomics Pipeline^55^ and PeptideDepot^56^. MS2 spectra were searched using the Mascot search engine against the complete UniProt proteome dataset (downloaded August 2019, 98,300 unique forward sequences) with a 0.1% FDR cutoff against a decoy database. The following parameters were used for peptide spectrum matching: trypsin (up to 2 missed cleavages), precursor mass tolerance of 7 ppm, 500 mmu fragment ion mass tolerance, phosphorylation as a variable modification (S/T/Y + 79.9963 Da), oxidation as a variable modification (M +15.9949), and SILAC labelling (K: +8.0142 Da, R: +10.0083 Da) and carbamidomethylation (C +57.0215) as a static modification.

Relative peptide abundance was determined using peak area integration and retention time alignment of selected ion chromatograms. The abundance of pY peptides was normalised using the spike in standard peptide LIEDpYTAK, and any samples without LIEDpYTAK sequenced were omitted. Statistical significance between the log2 mean abundance for each comparison was determined using Welch’s T test and corrected using the method of Storey^57^ as previously described. Post-translational modification signature enrichment analysis was performed as previously described^35^.

All post-PeptideDepot data processing was performed using Python 3.10 in the Windows Subsystem for Linux 2 in JupyterLab environments with the following dependencies: matplotlib (3.10.1), scipy (1.15.2), numpy (2.2.5), scikitlearn (1.6.1), pandas (2.2.3), and a custom ‘helpers’ module for improving matplotlib graph quality. The Holm Sidak algorithm for FWER correction was performed using a self-coded version of the algorithm as described by GraphPad Prism.

### 4.8 Data availability

All Western blot images, Floreada workspaces, and all code used for the analysis and presentation of our mass spectrometry data are publicly available on GitHub (https://github.com/Aurdeegz/Zap70-Itk-Inhibitor-Profiling). Mass spectrometry data are available from the ProteomeXchange Consortium via the PRIDE repository (Dataset ID: PXD069485).

## Acknowledgments

The authors would like to thank Dr. Xiaolei Su and his research team for providing the Jurkat CAR T cells for these experiments, as well as the group of Dr. Arthur Weiss for providing T2-Kb cells and Jurkat cells expressing the murine OT1 TCR & CD8. We would also like to wish Dr. Weiss a happy retirement and thank him for our years of enjoyable and productive collaborations. This work was supported by NIH grants P01AI091580 and 1S10OD036295.

## Contributions

Conceptualisation: AC; Methodology: AC; Investigation: AC, SST, AM, TR; Visualisation: AC; Funding acquisition: ARS; Project administration: AC; Supervision: AC, ARS; Writing: original draft: AC; Writing: review & editing: AC, ARS

## Additional information

The authors declare no competing interests. Generative AI was not used in the writing process, nor in the generation of figures for the manuscript.

## Notes

https://github.com/Aurdeegz/Zap70-Itk-Inhibitor-Profiling

### Competing Interest Statement

The authors have declared no competing interest.

