## Supporting Figures for "Phospotyrosine proteomics reveals novel Zap70 and Itk pathway targets downstream of TCR and CAR in Jurkat T cells"

**Abstract**: (200 words max)

ζ-associated protein of 70 kDa (Zap70) and interleukin 2 inducible T cell kinase (Itk) propagate the primary and CD28 integrated phosphotyrosine (pY) signalling, respectively, to achieve full T cell activation. Despite their canonical roles in T cell activation, our understanding of how each kinase controls canonical and noncanonical pY signalling is incomplete. Here, using three T cell activation methods (soluble antibodies, APC-pMHC/TCR, and CD19-CAR/Raji), we evaluated the two novel inhibitors RDN2150 (RDN, Zap70) and Soquelitinib (Soq, Itk) for their effects on T cell activation. We validated the published working concentrations of RDN and Soq on phosphorylation of key T cell signalling proteins and on T cell activation markers, finding that RDN provides more complete inhibition of T cell signalling and activation. We used LC-MS/MS to evaluate how RDN and Soq treatment affected the phosphotyrosine (pY) signalling and proteome of T cells, finding that RDN, as opposed to Soq, completely downregulated the TCR signalling pathway. Finally, we identified new, noncanonical pY sites responsive to RDN and Soq, providing new insight into the pathways regulated by Zap70 and Itk. Together, our work provides a basis for further study on RDN and Soq and a molecular roadmap for the effects of these inhibitors.

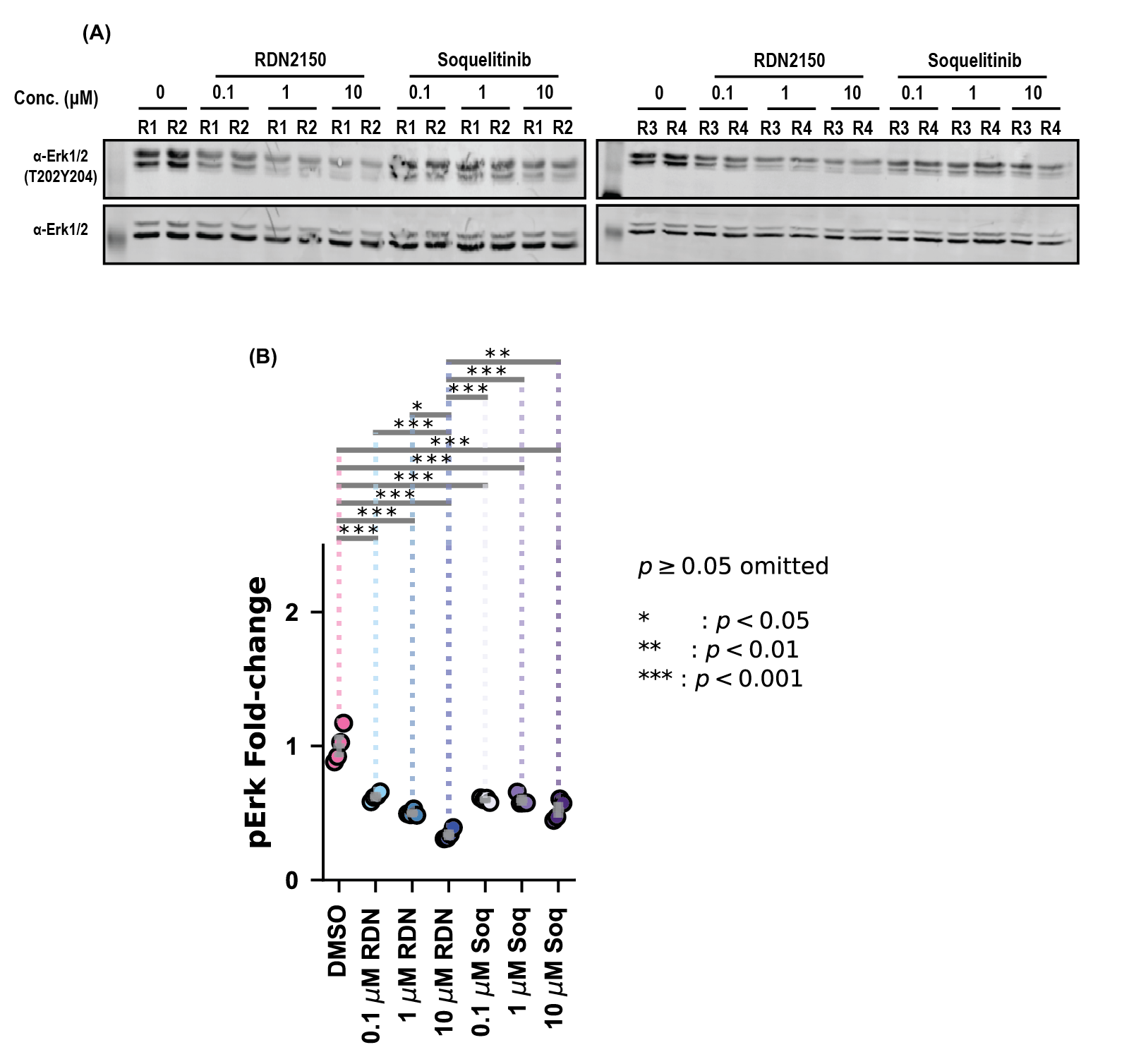

Supporting Figure 1: Treatment with low-levels of RDN2150 or Soquelitinib reduces Erk1/2 phosphorylation in unstimulated Jurkat T cells. (A) Western blot analysis of Erk1T202Y204/Erk2T185Y187 in unstimulated Jurkat T cells with (B) accompanying quantification. Statistical analysis for Western blots was performed using the Holm-Sidak correction of Fisher’s LSD pairwise comparisons.

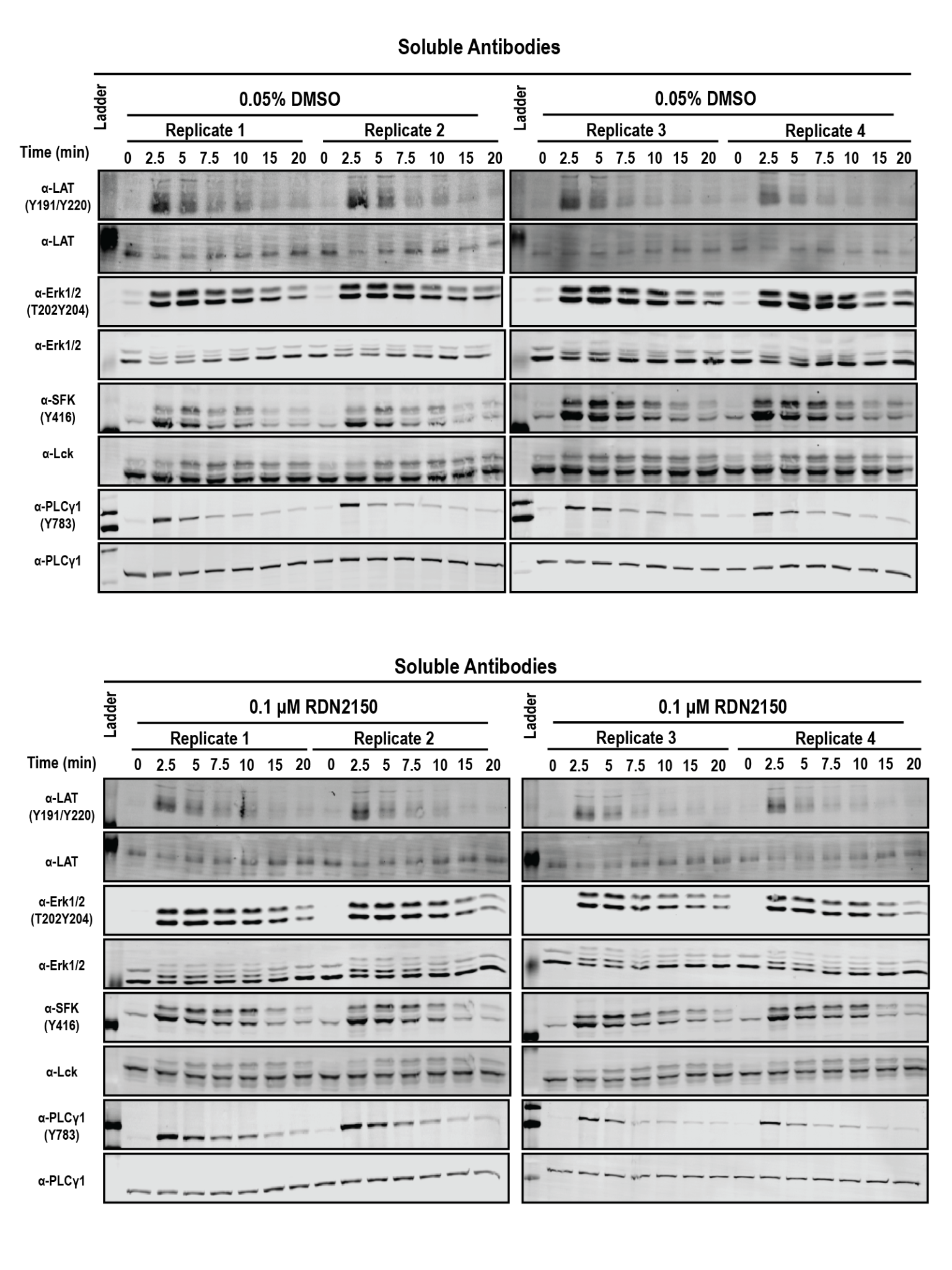

Supporting Figure 2: Legend on Page 6

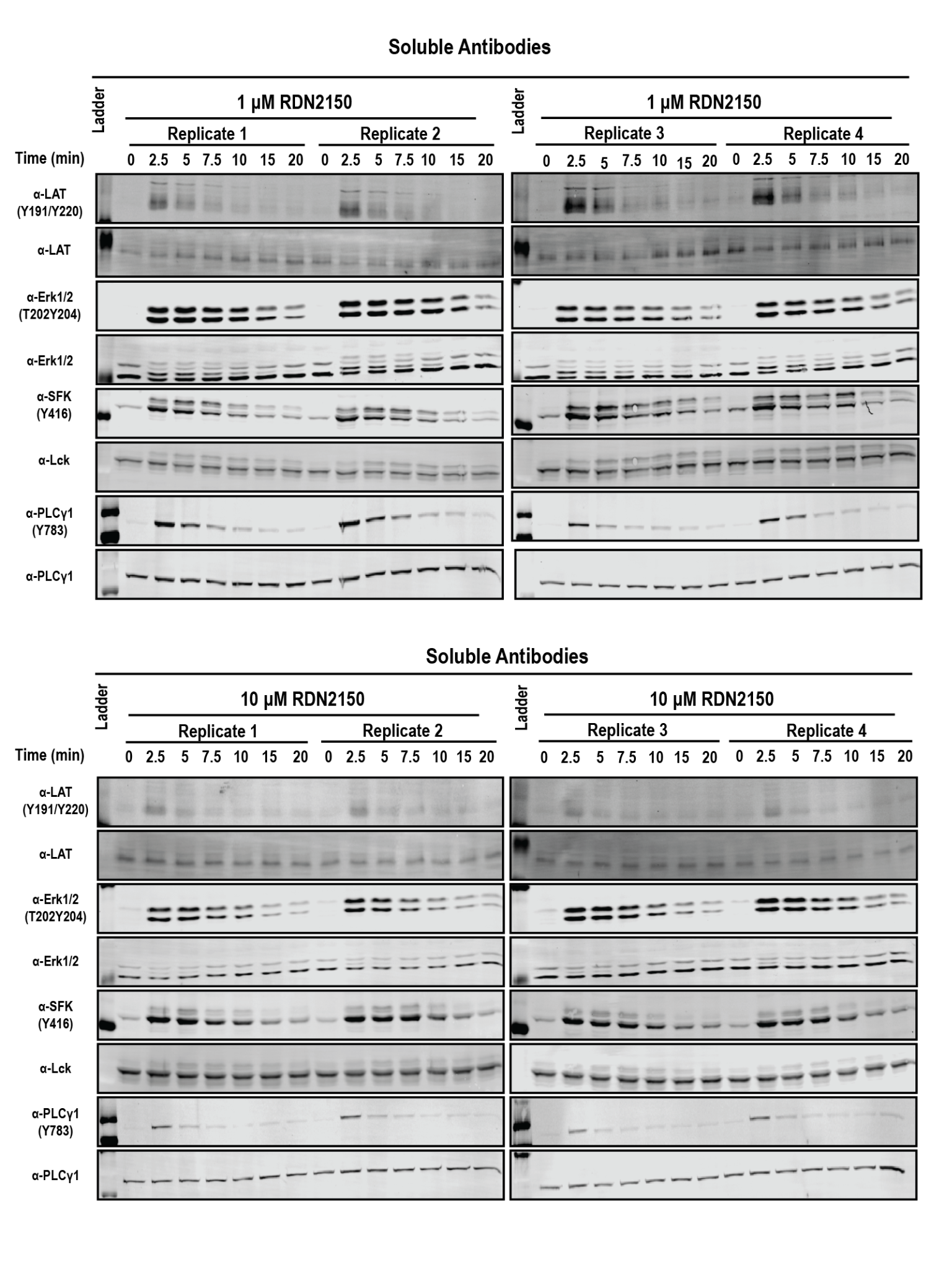

Supporting Figure 2: Legend on Page 6

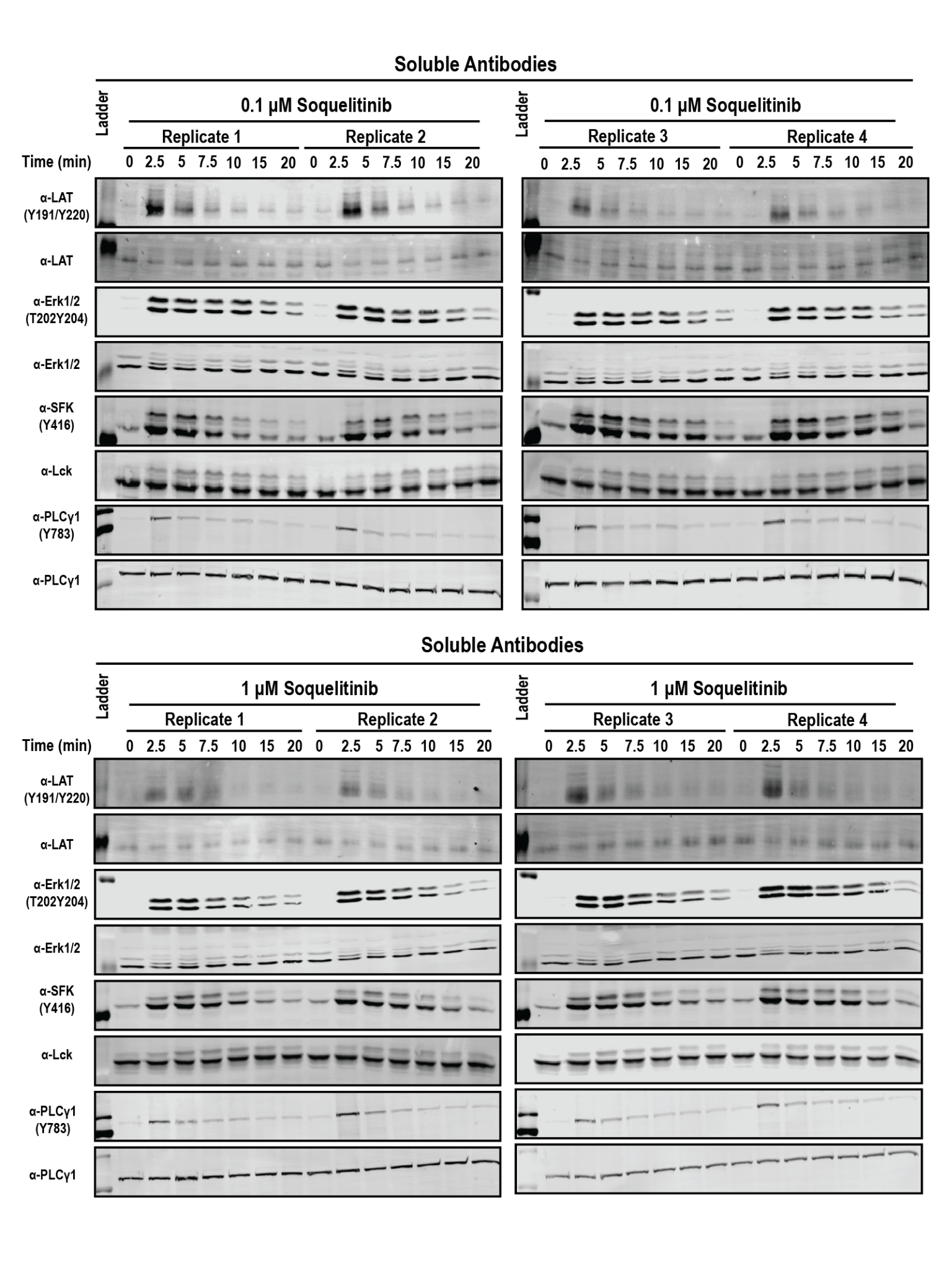

Supporting Figure 2: Legend on Page 6

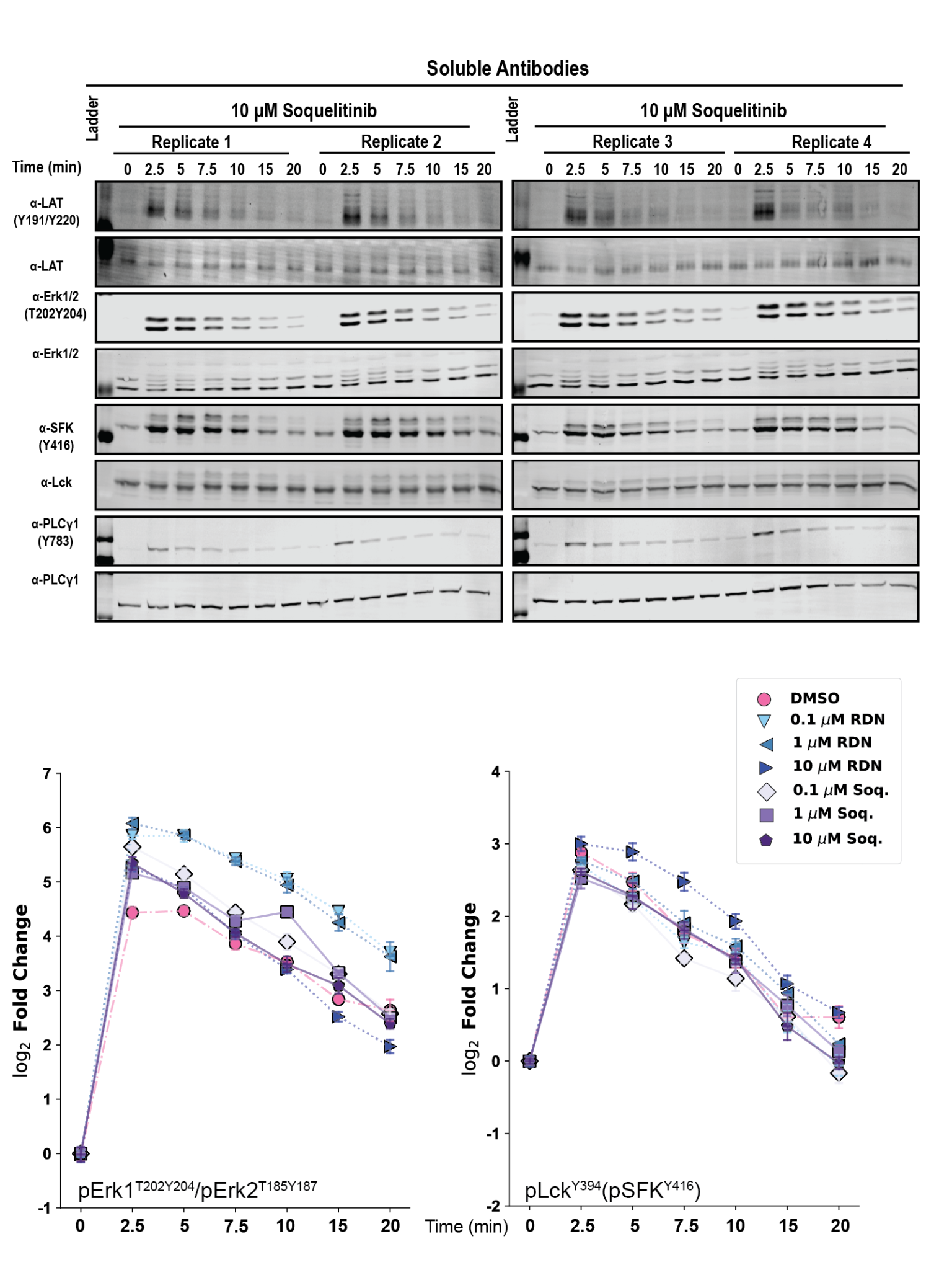

Supporting Figure 2 (Pages 3-6): Quantitative Western blots after time course soluble antibody crosslinking of CD3ε/CD28 and a titration of RDN2150/Soquelitinib reveals a dramatic increase in pY abundance of the key TCR phosphorylation sites LAT^Y220^, Erk1^T202Y204^/Erk2^T185Y187^, Lck^Y394^/SFK^Y416^, and PLCγ1^Y783^. Quantification for PLCγ1^Y783^ and LAT^Y220^ are shown in Figure 1. Statistical analysis for Western blots was performed using the Holm-Sidak correction of Fisher’s LSD pairwise comparisons and is available in Supporting Tables 3 and 4.

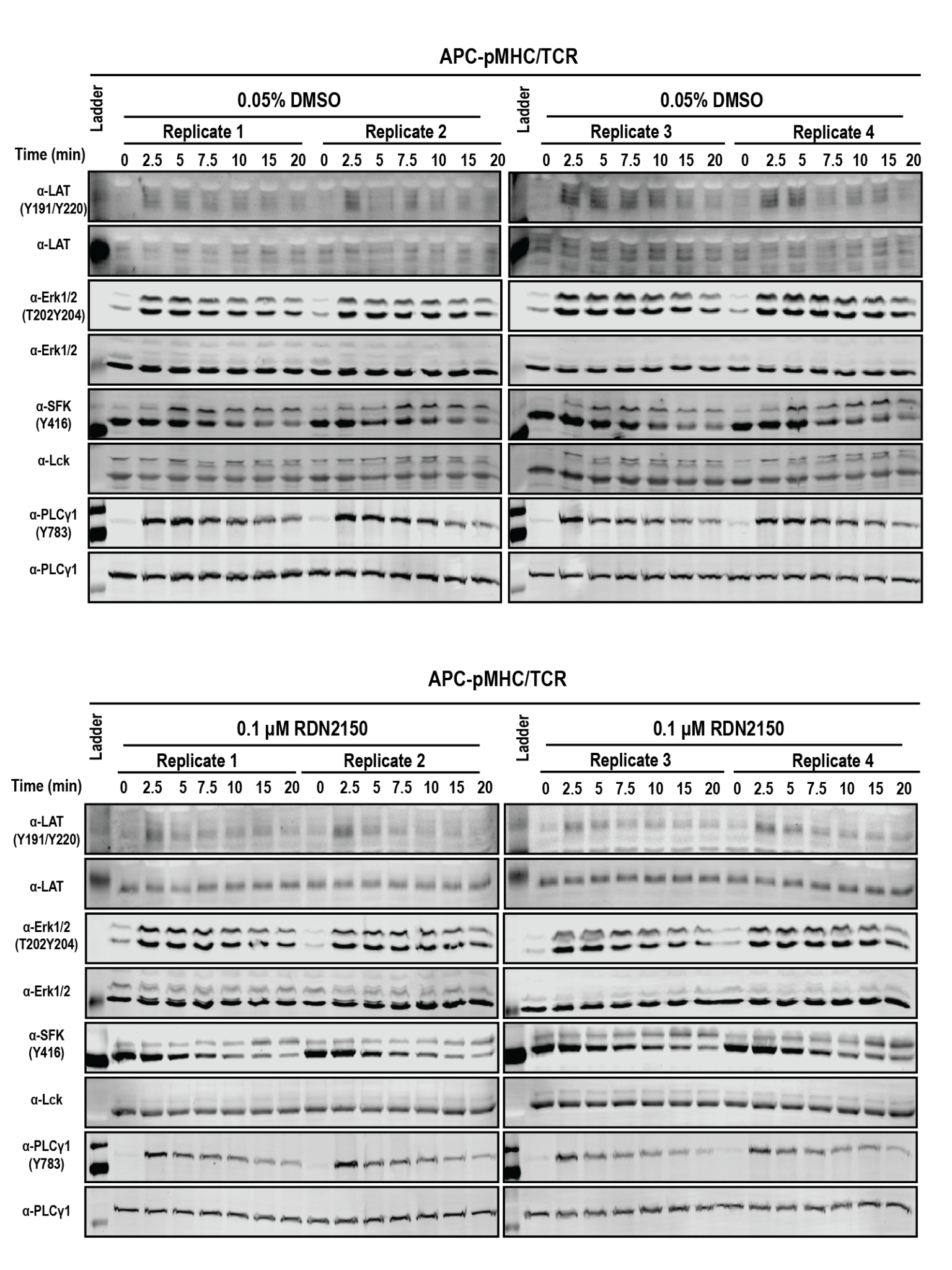

Supporting Figure 3: Legend on Page 11

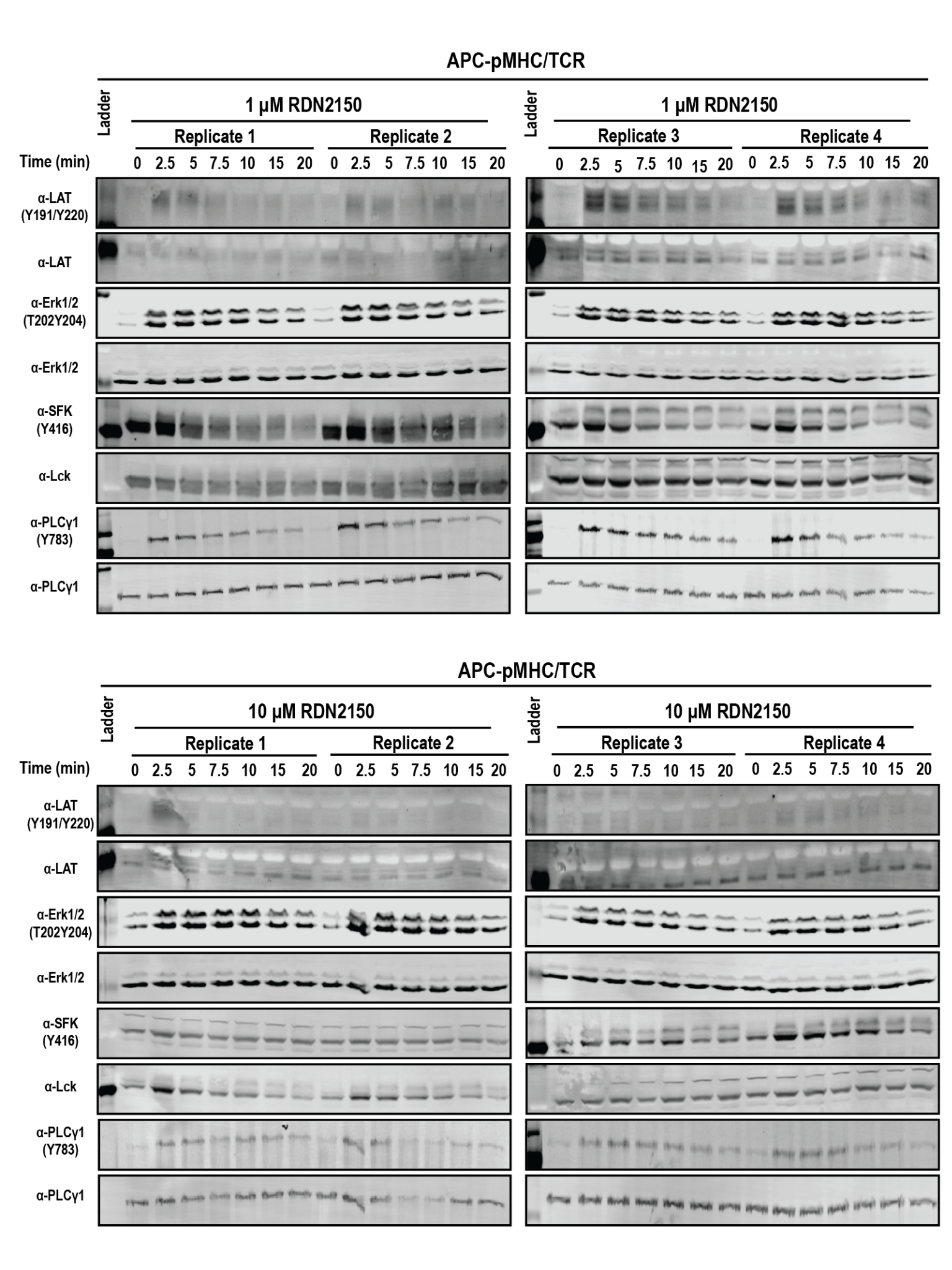

Supporting Figure 3: Legend on Page 11

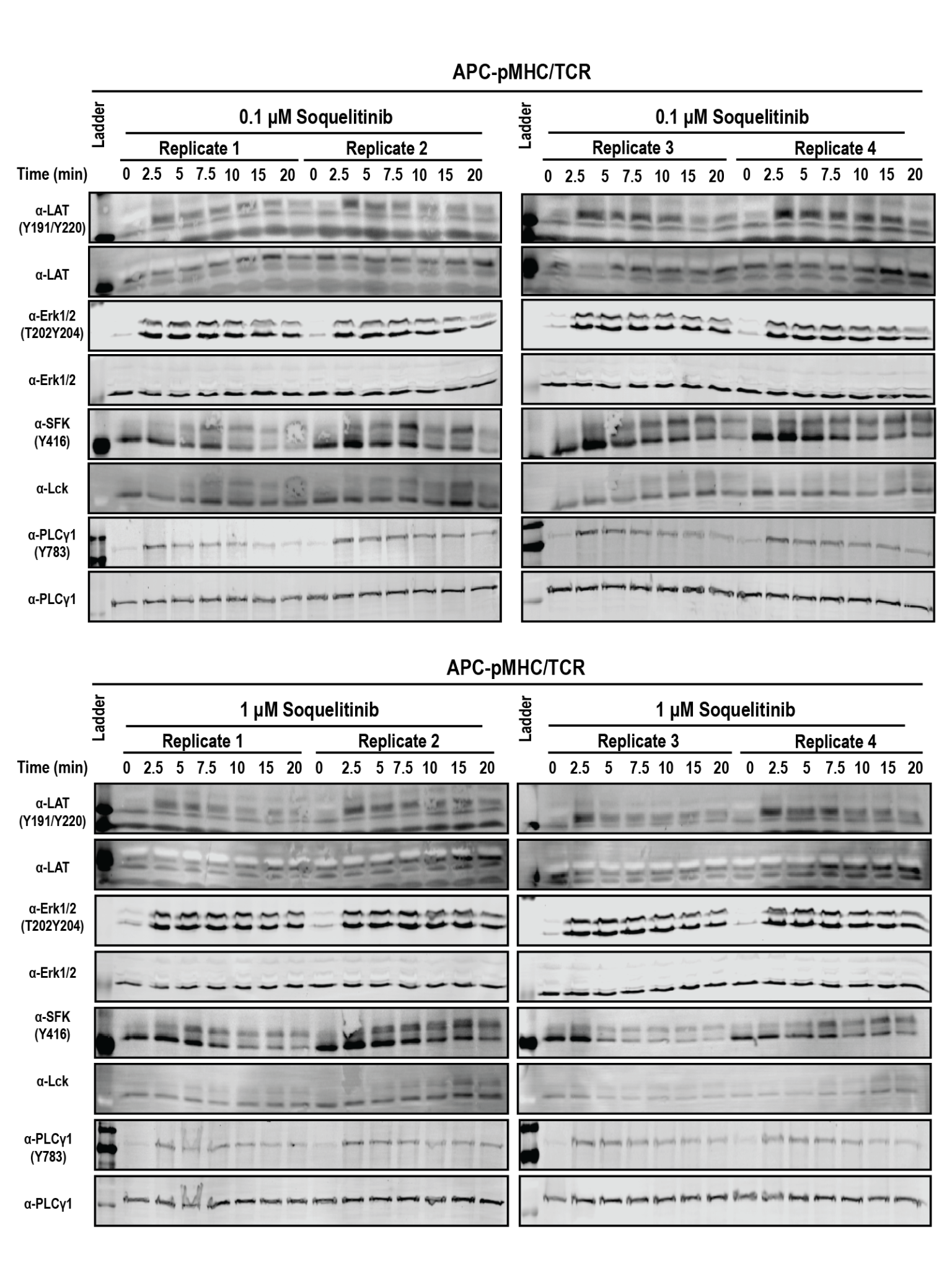

Supporting Figure 3: Legend on Page 11

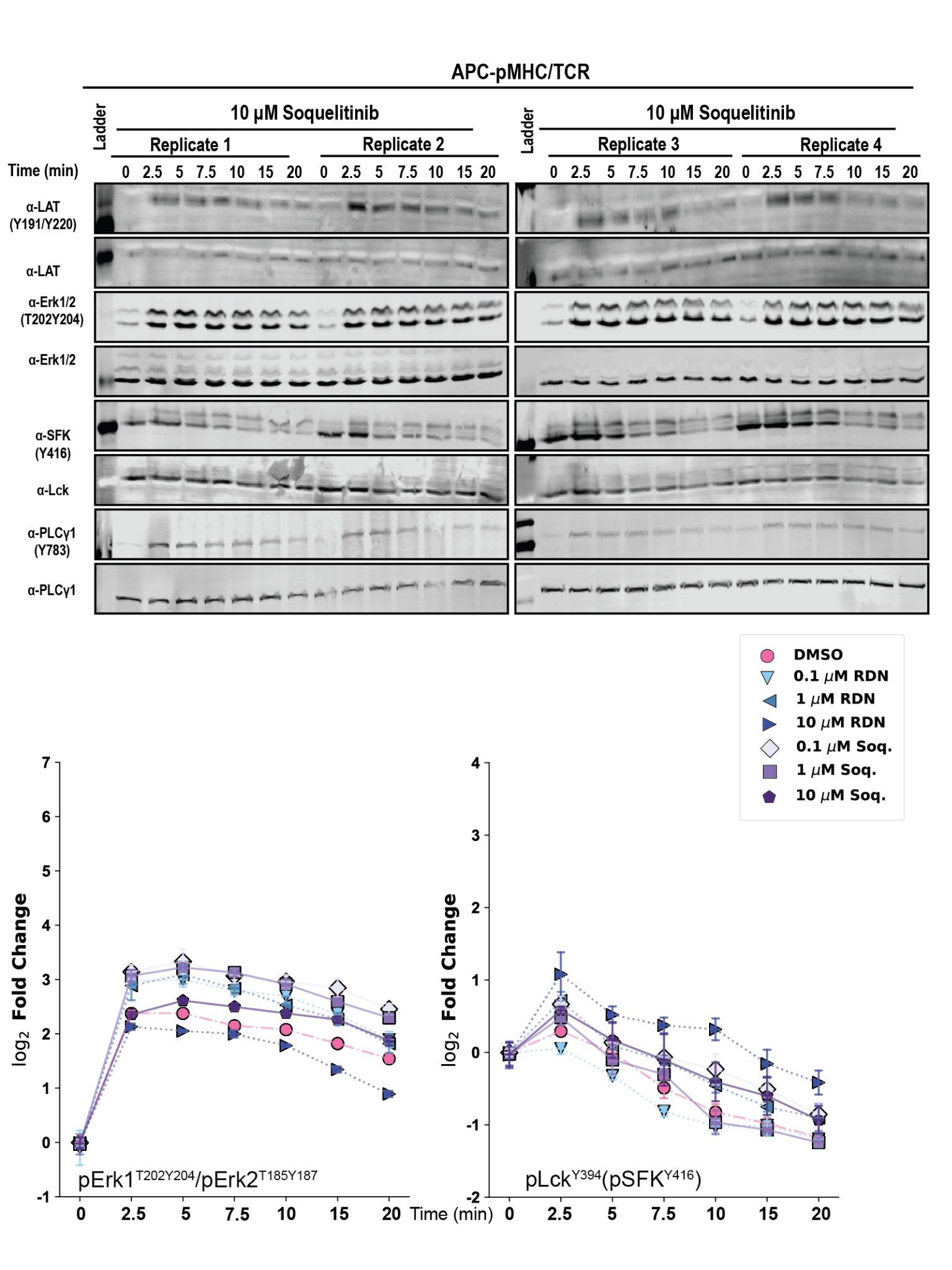

Supporting Figure 3 (Pages 8-11): Quantitative Western blots after time course co-culture of peptide-loaded T2-K^b^ cells with J^OT1^ cells and a titration of RDN2150/Soquelitinib reveals a modest increase in pY abundance of the key TCR phosphorylation sites LAT^Y220^, Erk1^T202Y204^/Erk2^T185Y187^, Lck^Y394^/SFK^Y416^, and PLCγ1^Y783^. Quantification for PLCγ1^Y783^ and LAT^Y220^ are shown in Figure 1. Statistical analysis for Western blots was performed using the Holm-Sidak correction of Fisher’s LSD pairwise comparisons and is available in Supporting Tables 7 and 8.

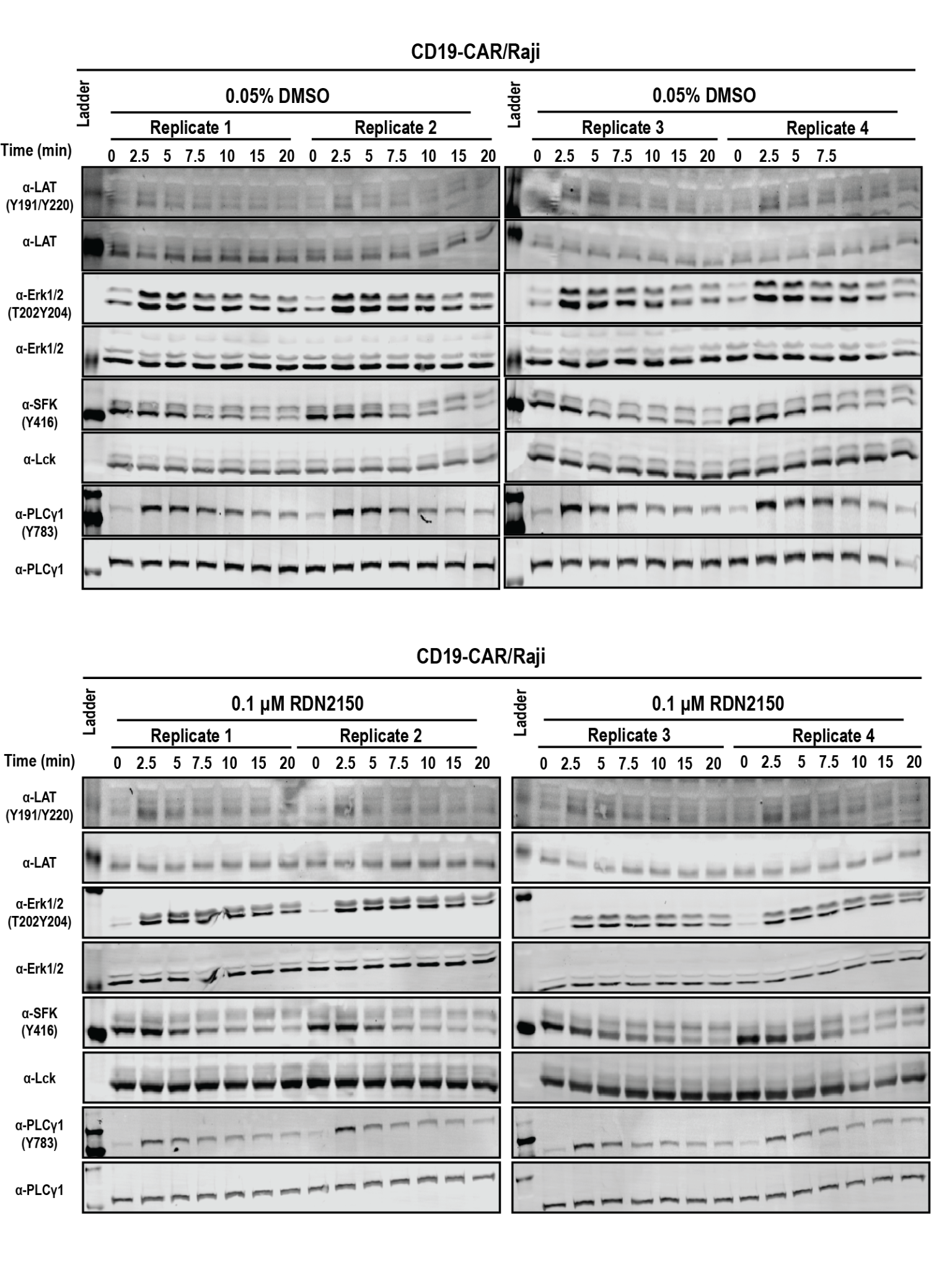

Supporting Figure 4: Legend on Page 16

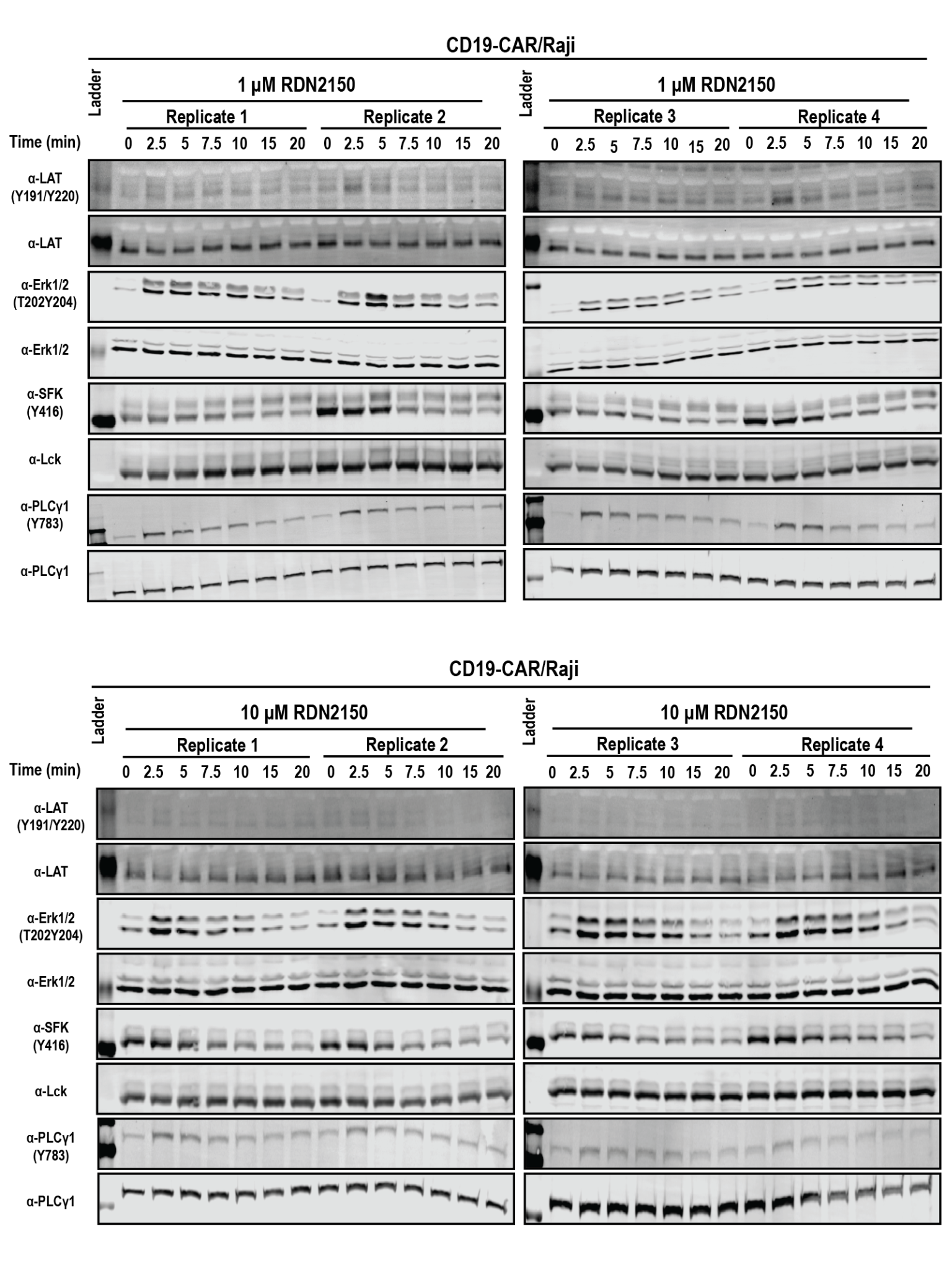

Supporting Figure 4: Legend on Page 16

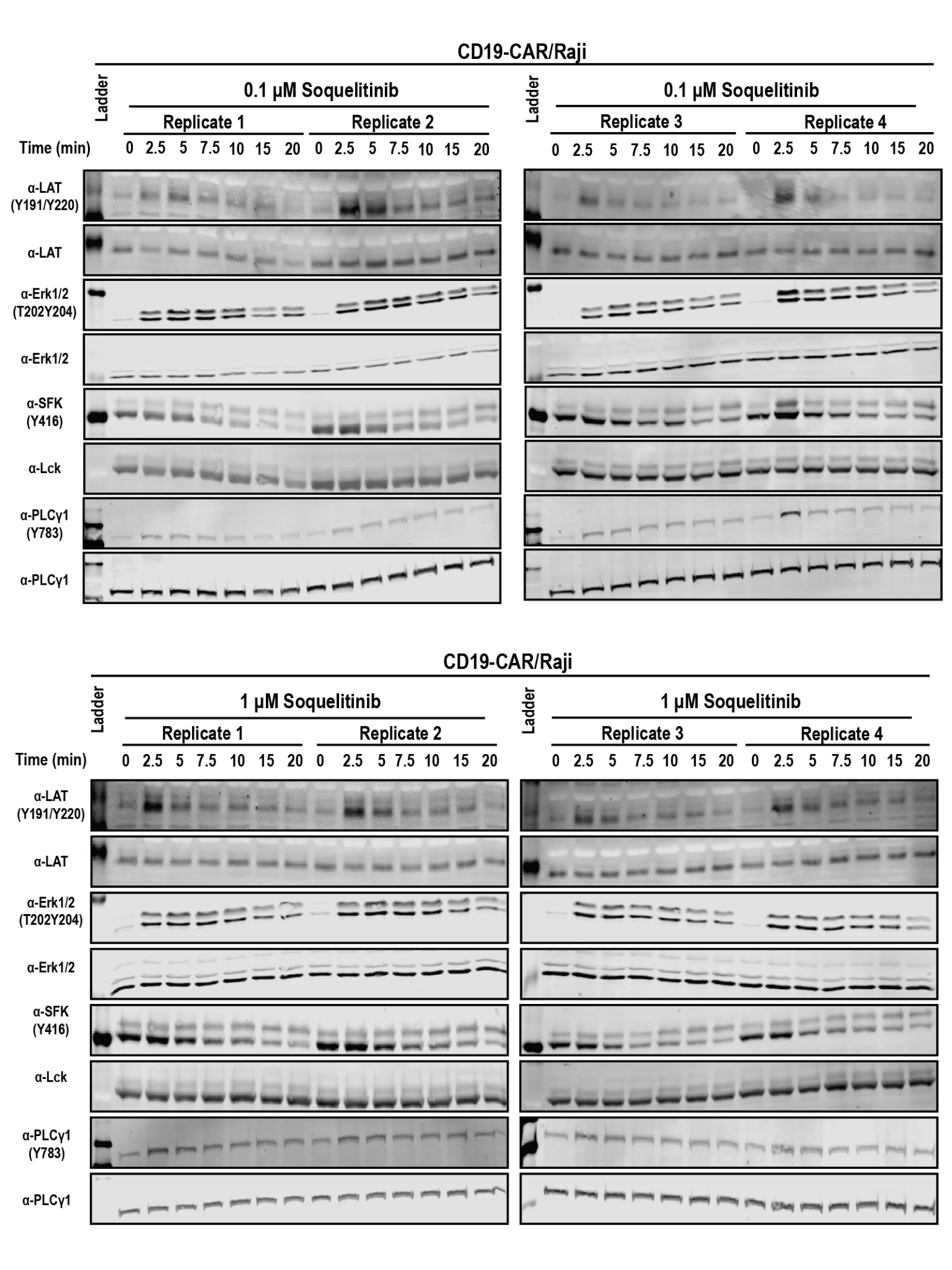

Supporting Figure 4: Legend on Page 16

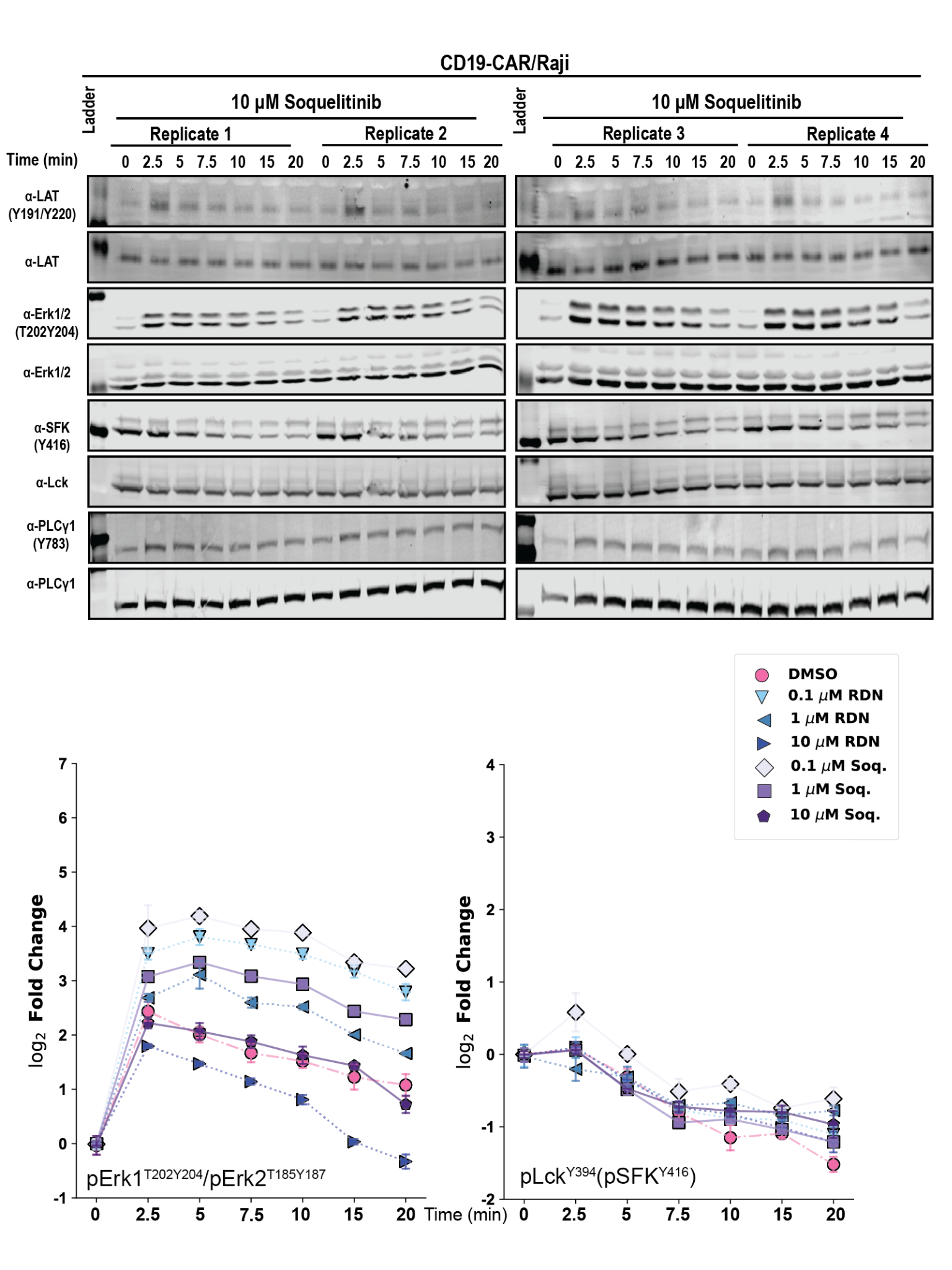

Supporting Figure 4 (Pages 13-16): Quantitative Western blots after time course co-culture of CD19-expressing Raji B cells with CD19-28ζ-CAR T cells and a titration of RDN2150/Soquelitinib reveals generally weak changes in pY abundance of the key TCR phosphorylation sites LAT^Y220^, Erk1^T202Y204^/Erk2^T185Y187^, Lck^Y394^/SFK^Y416^, and PLCγ1^Y783^. Quantification for PLCγ1^Y783^ and LAT^Y220^ are shown in Figure 1. Statistical analysis for Western blots was performed using the Holm-Sidak correction of Fisher’s LSD pairwise comparisons and is available in Supporting Tables 11 and 12.

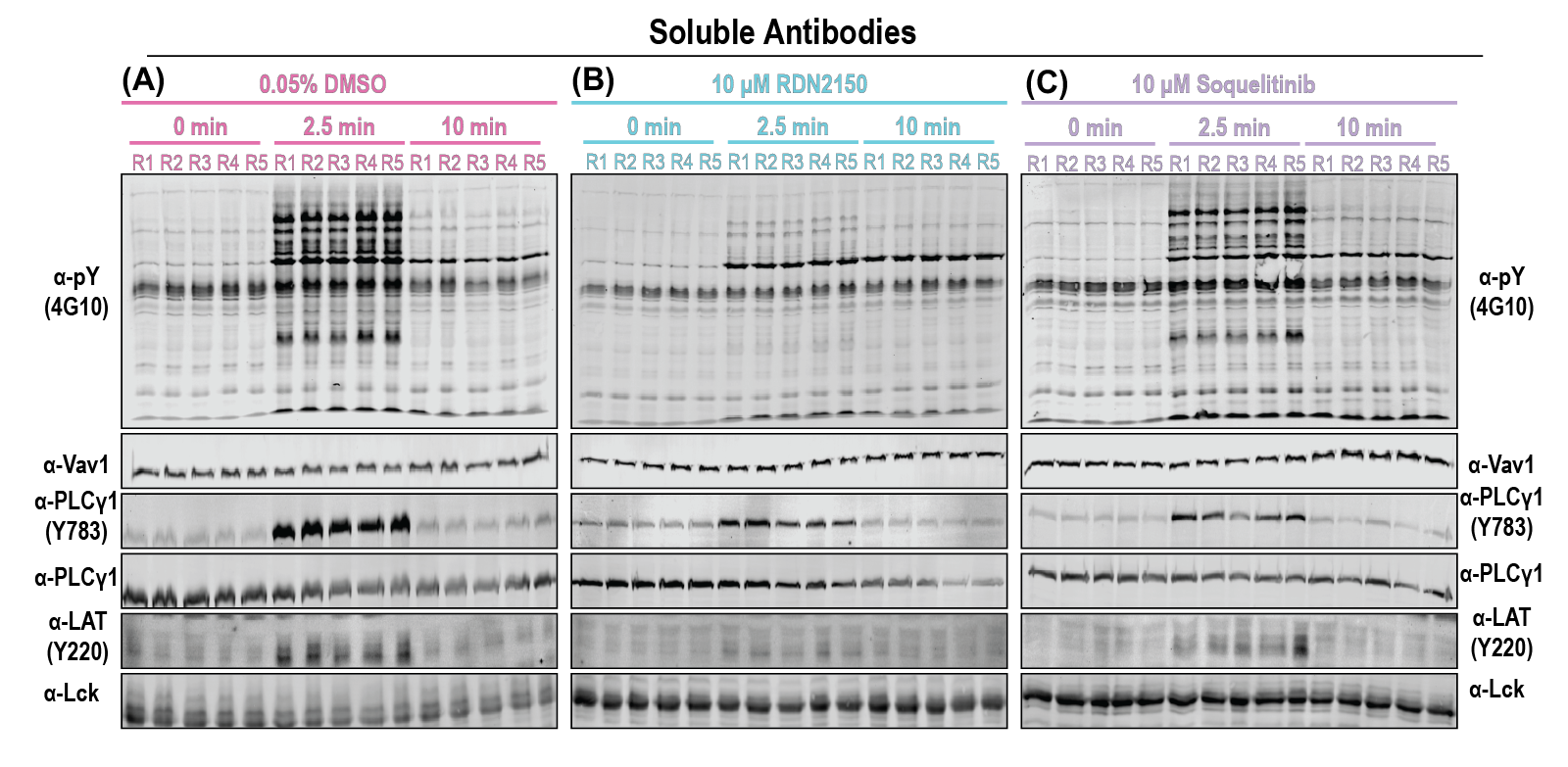

Supporting Figure 5: Western blot analysis of soluble antibody stimulation pY proteomics samples targeting: phosphotyrosine (pY, clone 4G10), Vav1 (loading control), PLCγ1^Y783^/PLCγ1, and LAT^Y220^/Lck in (A) DMSO control samples, (B) 10 μM RDN2150 samples, and (C) 10 μM Soquelitinib samples.

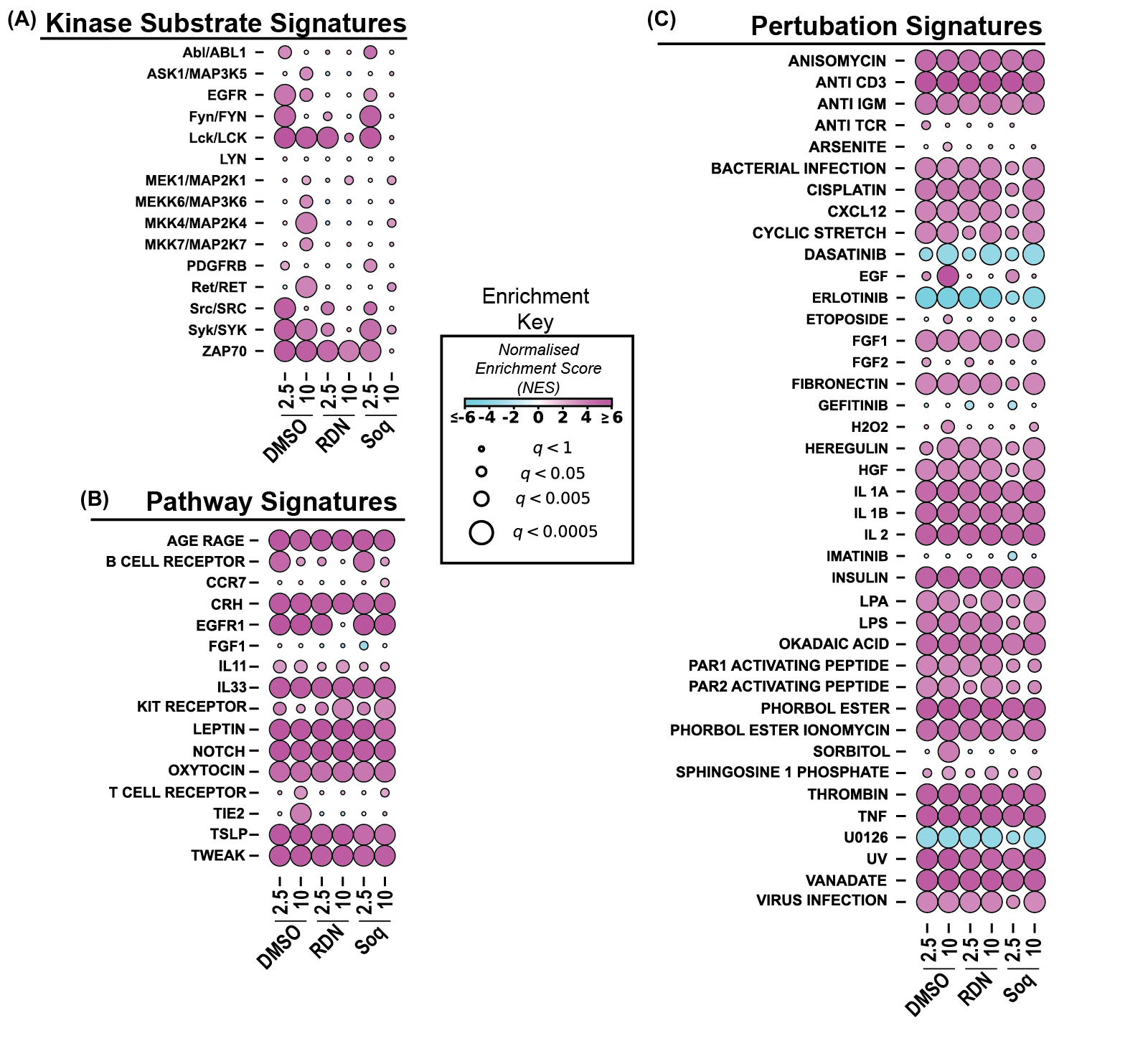

Supporting Figure 6: Post translational signature enrichment analysis (PTM-SEA) reveals soluble antibody stimulation induces pY signatures associated with increased kinase activity, signalling pathway engagement, and activating perturbations. (A) PTM-SEA on stimulation time course comparisons for PhosphoSitePlus Kinase Substrate Signatures. (B) PTM-SEA on stimulation time course comparisons for NetPath Pathway Signatures. (C) PTM-SEA on stimulation time course comparisons for PhosphoSitePlus Perturbation Signatures.

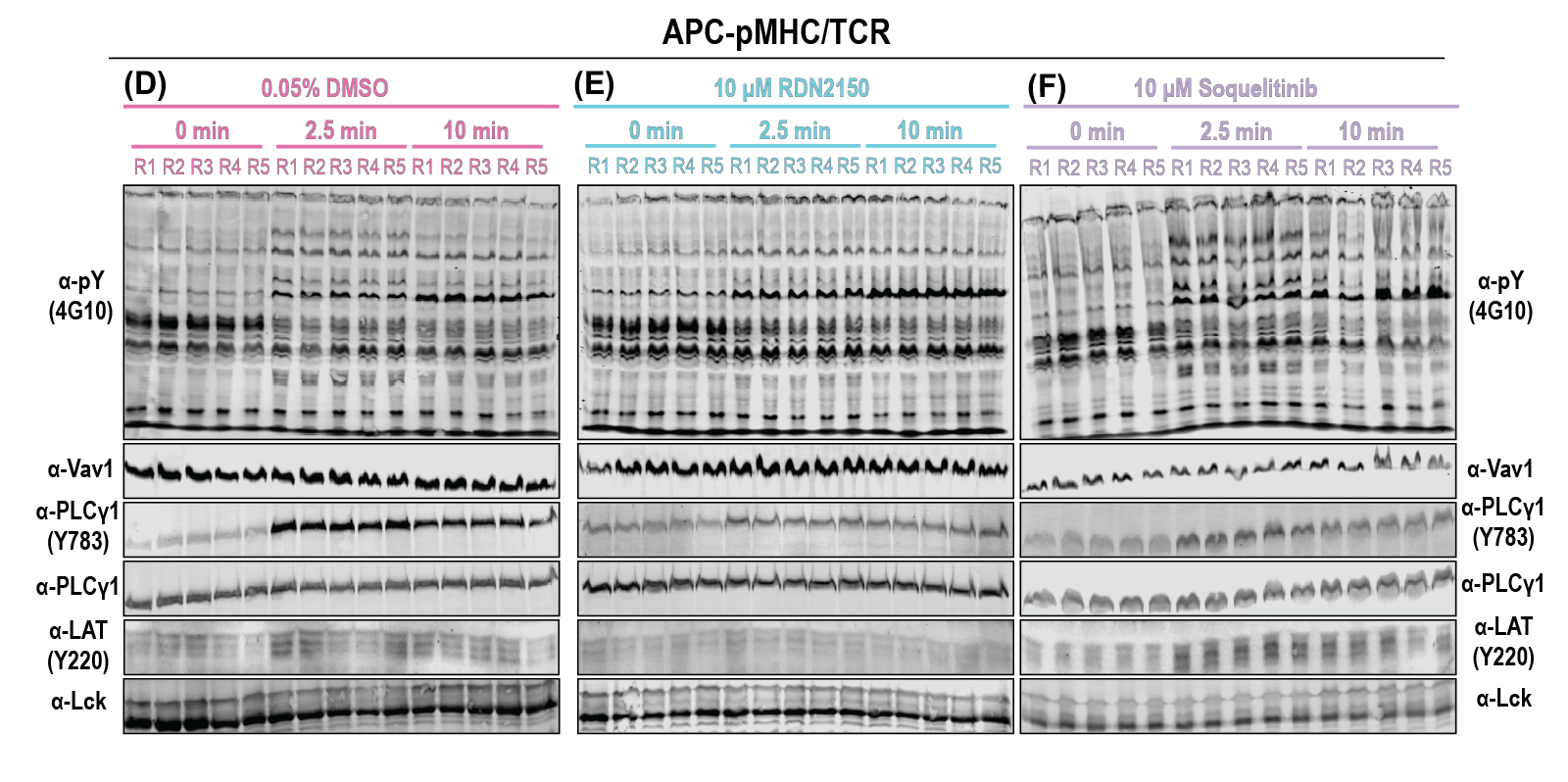

Supporting Figure 7: Western blot analysis of pMHC/TCR stimulation pY proteomics samples targeting: phosphotyrosine (pY, clone 4G10), Vav1 (loading control), PLCγ1^Y783^/PLCγ1, and LAT^Y220^/Lck in (A) DMSO control samples, (B) 10 μM RDN2150 samples, and (C) 10 μM Soquelitinib samples.

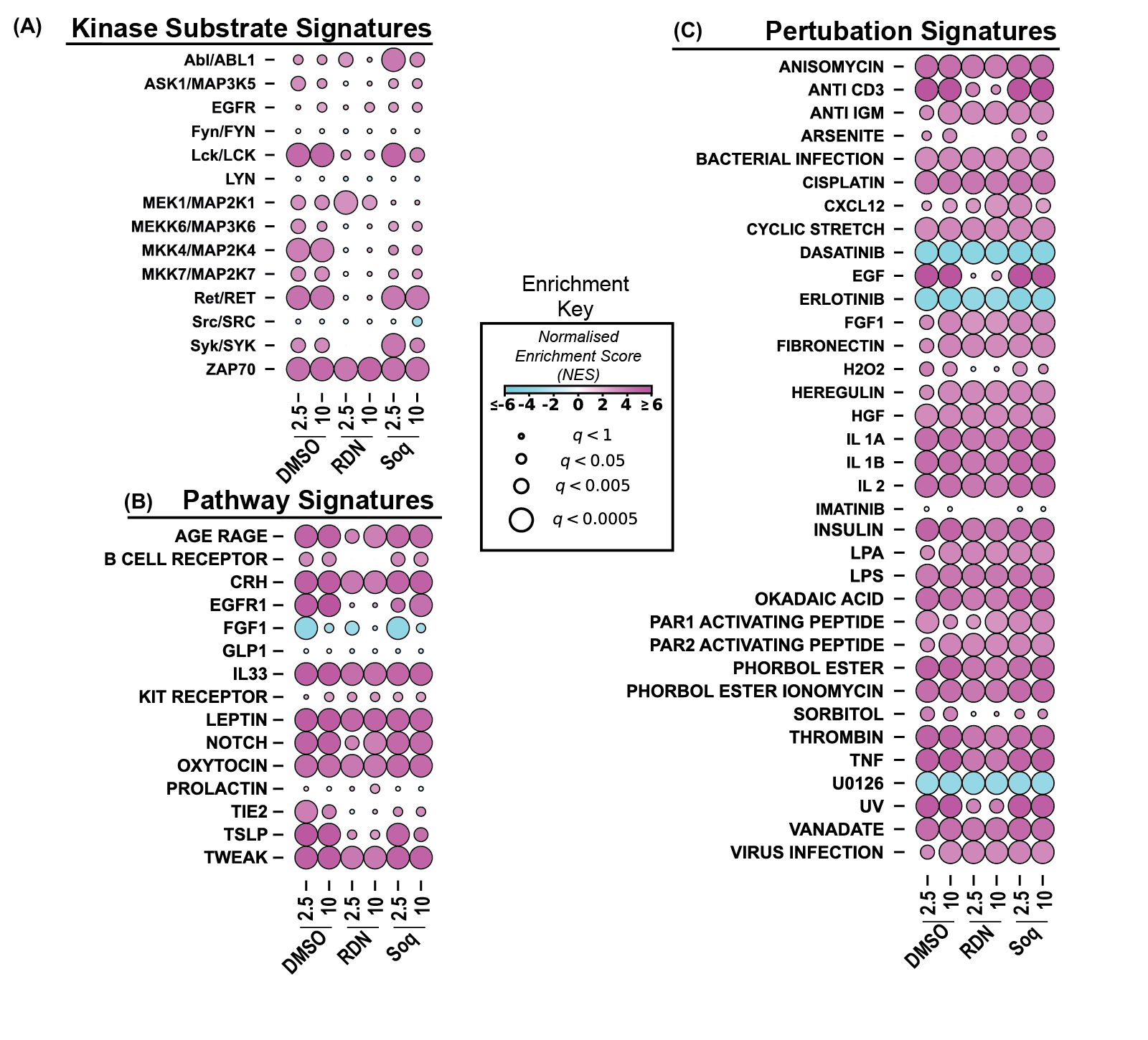

Supporting Figure 8: Post translational signature enrichment analysis (PTM-SEA) reveals pMHC/TCR stimulation induces pY signatures associated with increased kinase activity, signalling pathway engagement, and activating perturbations. (A) PTM-SEA on stimulation time course comparisons for PhosphoSitePlus Kinase Substrate Signatures. (B) PTM-SEA on stimulation time course comparisons for NetPath Pathway Signatures. (C) PTM-SEA on stimulation time course comparisons for PhosphoSitePlus Perturbation Signatures.

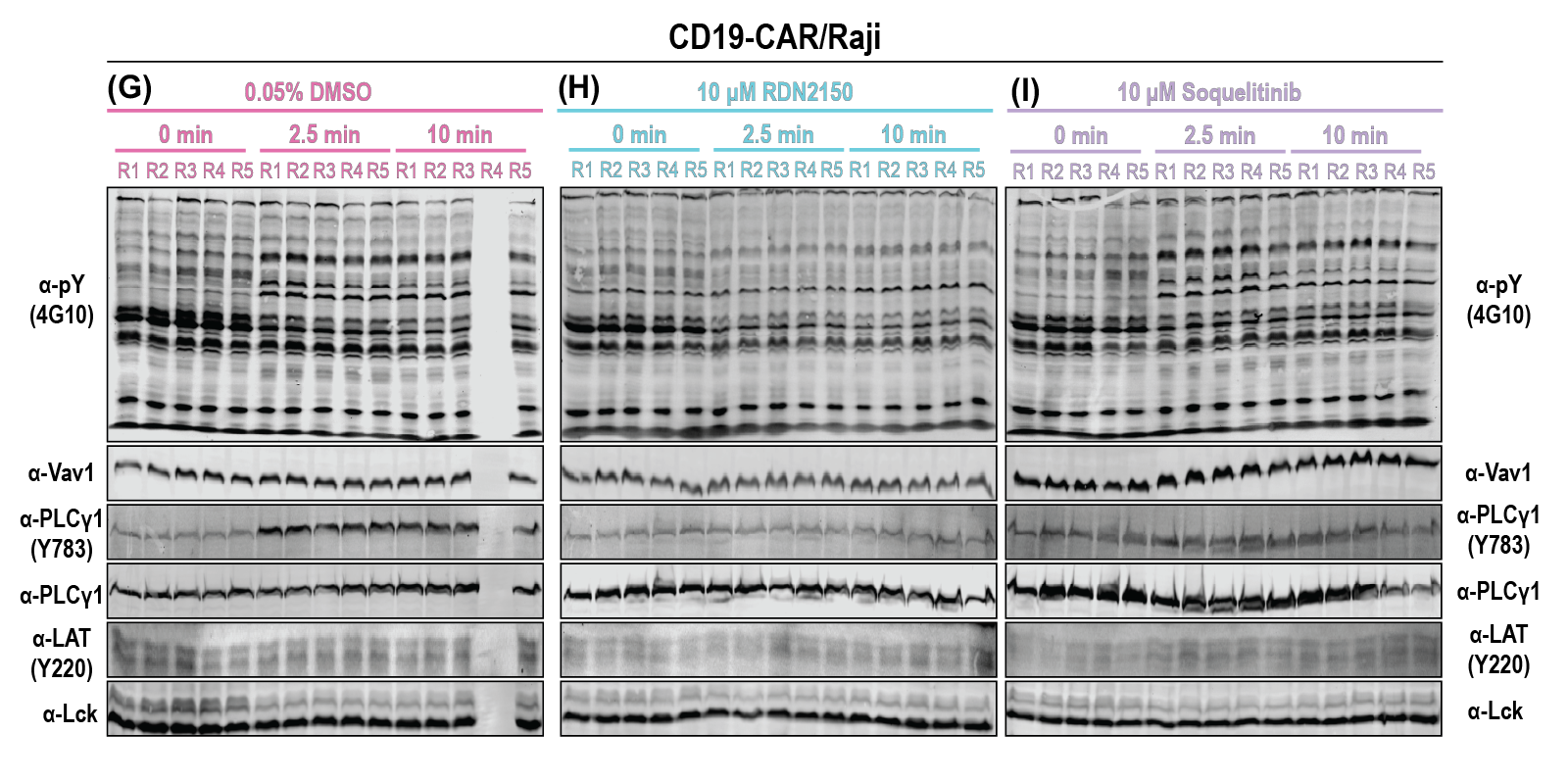

Supporting Figure 9: Western blot analysis of CAR/TAA stimulation pY proteomics samples targeting: phosphotyrosine (pY, clone 4G10), Vav1 (loading control), PLCγ1^Y783^/PLCγ1, and LAT^Y220^/Lck in (A) DMSO control samples, (B) 10 μM RDN2150 samples, and (C) 10 μM Soquelitinib samples.

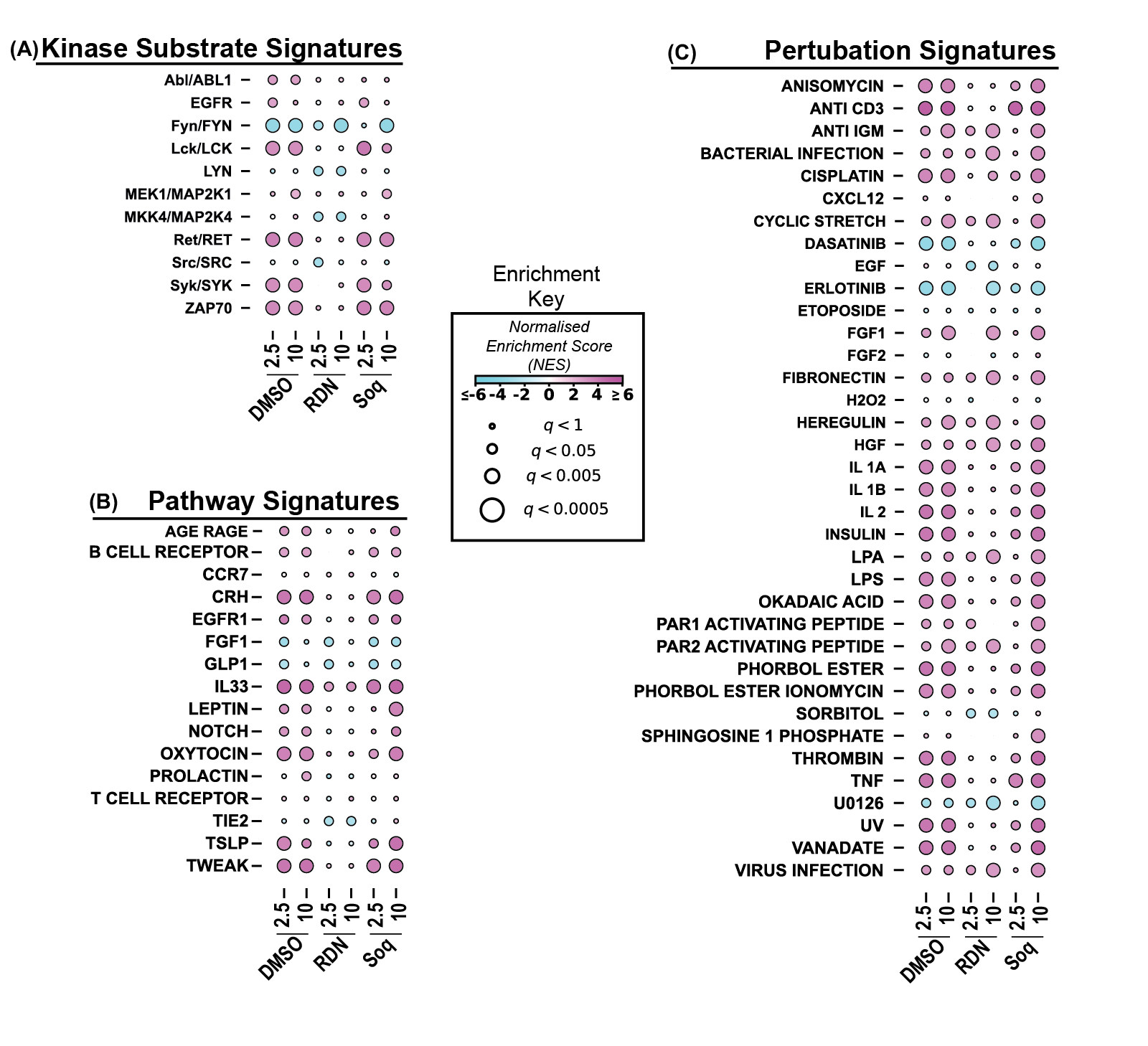

Supporting Figure 10: Post translational signature enrichment analysis (PTM-SEA) reveals CAR/TAA stimulation weakly engages pY signatures associated with increased kinase activity, signalling pathway engagement, and activating perturbations. (A) PTM-SEA on stimulation time course comparisons for PhosphoSitePlus Kinase Substrate Signatures. (B) PTM-SEA on stimulation time course comparisons for NetPath Pathway Signatures. (C) PTM-SEA on stimulation time course comparisons for PhosphoSitePlus Perturbation Signatures.

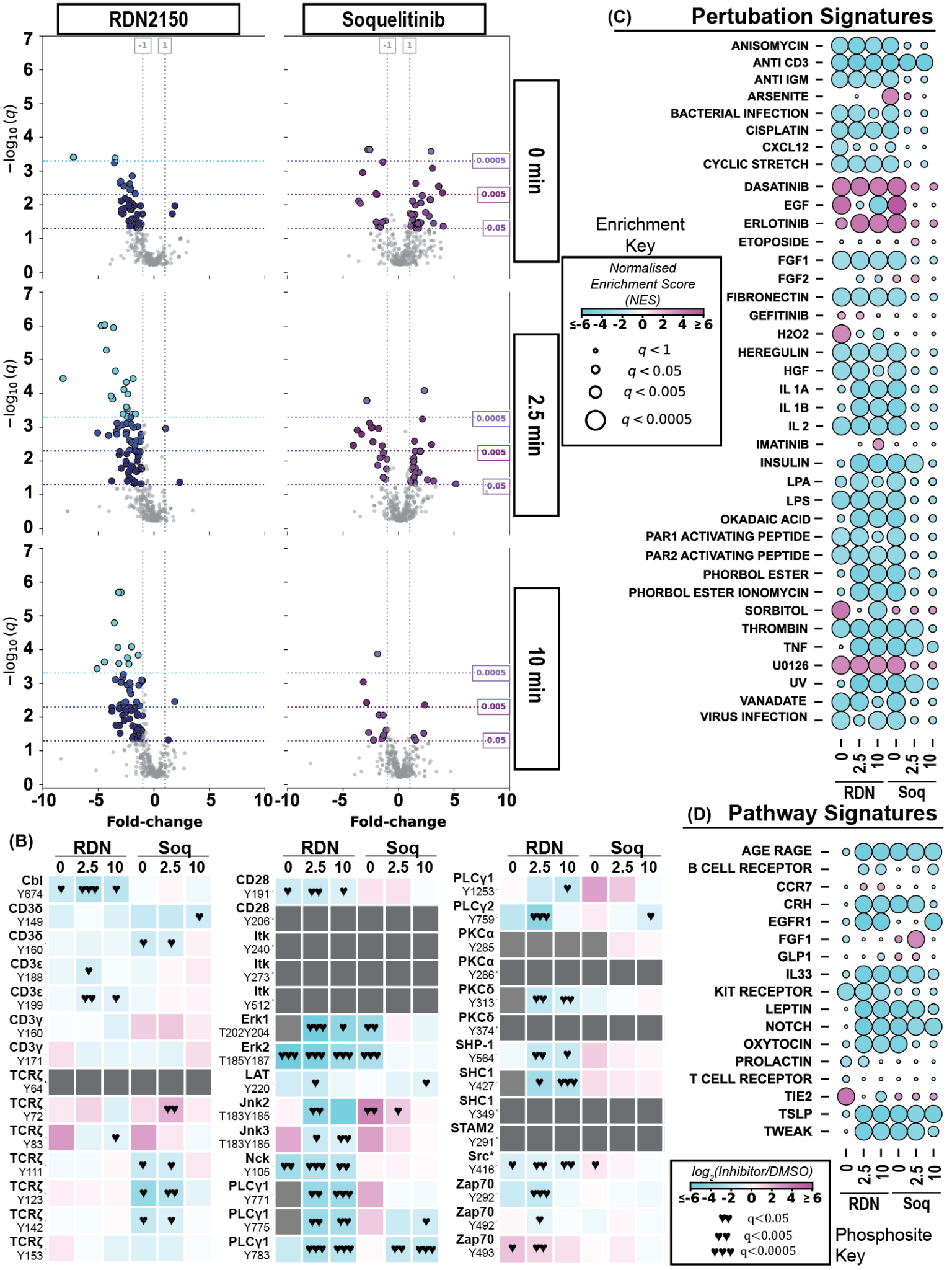

Supporting Figure 11: 10 RDN2150 treatment downregulates global pY abundance and the TCR signalling pathway more than Soquelitinib treatment. (A) Volcano plot analysis of pY proteomics data comparing RDN2150 and Soquelitinib treatment to a DMSO control in pMHC/TCR stimulated samples for 0 (top row), 2.5 (middle row), and 10 (bottom row) minutes. (B) Heat maps comparing RDN2150 and Soquelitinib treatment to a DMSO control for specific pY sites in the T cell signalling pathway. ♥ indicates q < 0.05, and ♥♥ indicates q < 0.005, and ♥♥♥ indicates q < 0.0005. (C-D) Post-translational modification signature enrichment analysis (PTM-SEA) on DMSO comparison data showing PhosphoSitePlus Perturbation Signatures and NetPath Pathway Signatures, respectively.

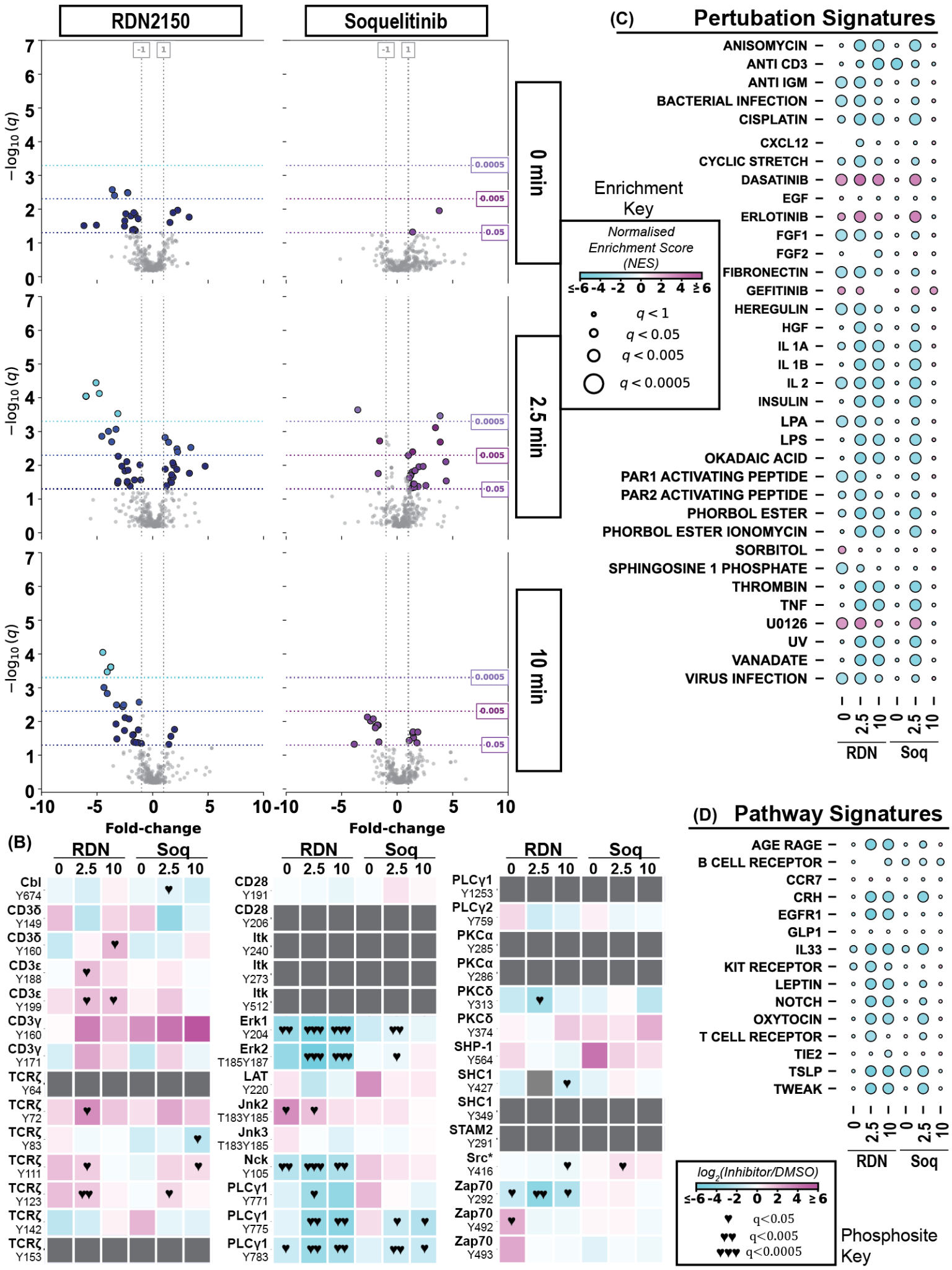

Supporting Figure 12: RDN2150 treatment downregulates global pY abundance and the CAR signalling pathway more than Soquelitinib treatment. (A) Volcano plot analysis of pY proteomics data comparing RDN2150 and Soquelitinib treatment to a DMSO control in CAR/TAA stimulated samples for 0 (top row), 2.5 (middle row), and 10 (bottom row) minutes. (B) Heat maps comparing RDN2150 and Soquelitinib treatment to a DMSO control for specific pY sites in the T cell signalling pathway. ♥ indicates q < 0.05, and ♥♥ indicates q < 0.005, and ♥♥♥ indicates q < 0.0005. (C-D) Post-translational modification signature enrichment analysis (PTM-SEA) on DMSO comparison data showing PhosphoSitePlus Perturbation Signatures and NetPath Pathway Signatures, respectively.
